# Essential role of NLRC5 in cancer immune surveillance and cancer immunoediting

**DOI:** 10.1101/2025.02.03.636144

**Authors:** Akhil Shukla, Anny Armas Cayarga, Jean-François Lucier, Madanraj Appiya Santharam, Dominique Lévesque, François-Michel Boisvert, Sheela Ramanathan, Subburaj Ilangumaran

## Abstract

A key mechanism of immune escape from CD8^+^ T cell-mediated tumor control occurs via downregulation of NLRC5, the IFNγ-induced transcriptional activator of MHC class-I. As NLRC5 deficiency does not abrogate CD8^+^ T cell development, we investigated whether NLRC5-dependent antitumor immune mechanisms are required for immune surveillance. Development of 3-methylcholanthrene (MCA)-induced endogenous fibrosarcoma was studied in *Nlrc5^-/-^* mice with *Nlrc5^+/+^* and *Rag1^-/-^* mice serving as controls. *Nlrc5^-/-^* and *Rag1^-/-^* mice showed increased propensity to develop MCA-induced tumors with elevated growth rate compared to *Nlrc5^+/+^* mice, and displayed significantly reduced survival. Tumors from *Nlrc5^+/+^* and *Nlrc5^-/-^*mice, but not from *Rag1^-/-^* mice, contained necrotic areas and displayed T cell infiltration. Tumor cell lines established from MCA-induced tumors were evaluated for their sensitivity to immune-mediated growth control following implantation into immunocompetent C57BL/6 and immunodeficient *Rag1^-/-^* hosts. Tumors formed by *Nlrc5^+/+^* tumor cell lines progressed unhindered in C57BL/6 hosts that reflected their immunoedited status, whereas cell lines from *Nlrc5^-/-^* and *Rag1^-/-^*tumors were efficiently controlled, indicating their non-immunoedited status. Proteomic analysis by mass spectrometry followed by pathway analysis revealed enrichment of granzyme-mediated cytolytic pathway in *Nlrc5^+/+^* tumors that were absent in *Nlrc5^-/-^* tumors, which showed enrichment of humoral and innate immune pathways. Overall, our findings show that NLRC5 is required for robust tumor immune surveillance and tumor immunoediting and that compensatory humoral and innate immune mechanisms activated by the loss of NLRC5 are insufficient for cancer immune surveillance and cancer immunoediting.

## Introduction

Immune evasion is a hallmark of cancer [1]. A key mechanism underlying the ability of cancer cells to evade killing by the adaptive immune system is the downmodulation of major histocompatibility class-I (MHC-I) molecules that present tumor antigenic peptides to CD8^+^ cytotoxic T lymphocytes (CTL) [2–5]. Reduced MHC-I expression often correlates with high tumor grading, disease progression, reduced patient survival and failure of CTL-based immunotherapies [4,6–9]. Loss of MHC-I can arise from deletions or mutations that give rise to hard/irreversible lesions, or from epigenetic repression that results in soft/reversible lesions [3,10–12]. The MHC-I antigen presentation pathway is linked to proteasomal cleavage of aged and misfolded proteins [13–15]. The peptides are transported across the endoplasmic reticulum by TAP1/TAP2 proteins, loaded on to MHC-I by TAPBP, and stable MHC-I:peptide complexes presented on the cell surface [16–18]. Thus, defective expression of any key component of the MHC-I antigen processing machinery (APM) can also result in impaired MHC-I expression [2,9,19–23]. The MHC-I:peptide complexes expressed on all nucleated cells are constantly scanned by CD8^+^ T lymphocytes, which detect and kill potentially neoplastic cells displaying tumor antigenic peptides, as a key mechanism of cancer immune surveillance. Cancer cells overcome this immune surveillance through immunoediting, which they achieve by downmodulating or mutating dominant tumor antigens and by reducing MHC-I expression [24,25].

NOD-Like Receptor CARD domain containing 5 (NLRC5) is a member of the NLR family proteins, which function as innate immune sensors of pathogen- or danger- associated molecular patterns [26,27]. NLRC5 attenuates LPS-induced NF-κB activation and reduces the induction of TNFα, IL-6 and IL-1β genes [28–32]. Similarities between NLRC5 and NLRA led to the discovery that NLRC5 transactivates MHC-I and APM genes [33,34]. These gene promoters contain *cis-*regulatory elements that bind transcription factors with which NLRC5 interacts and assembles a transcriptional enhanceosome [34–37]. NLRC5 is strongly induced by IFNγ [34]. Despite NLRC5 being the key transcriptional activator of MHC-I and the loss of β2M or MHC-I impairs CD8^+^ T cell development, NLRC5-deficient mice show only marginal reduction in CD8^+^ T cell numbers, possibly due to high basal MHC-I expression in thymic epithelial cells [38–45].

TCGA data from diverse cancers revealed that loss of NLRC5 expression is the most prevalent defect among MHC-I-related genes and correlates with reduced CTL infiltration, poor patient survival and unresponsiveness to immune checkpoint blockade therapy [46,47]. We and others have shown that stable expression of NLRC5 in MHC-I low cancer cells upregulates MHC-I and APM genes and promotes tumor antigen presentation and tumor control by CTLs [48,49]. Based on the above findings, NLRC5 is postulated to promote cancer immune surveillance [50], but this has not yet been experimentally validated. In this study we investigated whether NLRC5 is critical or dispensable for cancer immune surveillance and cancer immunoediting using mouse genetic models.

## Materials and Methods

### Mice

*Nlrc5^-/-^* mice and control *Nlrc5^+/+^* in C57BL/6J (Jax Strain #:000664) background have been previously described [51]. Functional impact of NLRC5 deficiency was confirmed by impaired upregulation of MHC-I and APM genes in splenocytes following IFNγ stimulation (Supplementary Figure S1). *Rag1^-/-^*mice in C57BL/6J background were purchased from the Jackson laboratory (B6.129S7-*Rag1^tm1Mom^*/J; Strain #:002216). All strains of mice were housed in the same breeding and experimentation rooms of the Université de Sherbrooke specific pathogen-free animal facility. Mice were housed in ventilated cages with 14/10 hours day/night cycle and fed with normal chow *ad libitum*. All experiments on mice were carried out during daytime with the approval of the Université de Sherbrooke Ethics Committee for Animal Care and Use (Protocol ID 2023-4043).

### Flow cytometry

Spleens from *Nlrc5^-/-^* and *Nlrc5^+/+^*mice were collected into sterile PBS containing 2% fetal calf serum in Petri dishes and crushed between glass slides to obtain single cell suspension. Red-blood cells were removed by incubating the cell pellet in 2 mL erythrocyte lysis buffer (150mM NH_4_Cl, 10mM KHCO_3_, 0.1mM Na_2_EDTA, pH 7.2-7.4) for 2 minutes and washed in PBS-FCS. Aliquots of cells were incubated with mouse IFNγ (20 ng mL^−1^) for 8 h and 24 h. Control and IFNγ stimulated cells were labelled for flow cytometry using fluorochrome conjugated antibodies listed in Table S1 as previously described {Kandhi, 2022 #675}. Cells were analyzed using CytoFLEX (Beckman Coulter, CA USA) flow cytometer and data analyzed using Flowjo software (BD Bioscience, OR, USA, version 10.9.0). MHC-I expression was evaluated using an antibody that detects H-2K^b^/D^b^ on CD45 gated CD3+CD4+, CD3+CD8+ and CD3-B220+ cells (Table S1) and expressed as mean fluorescence intensity.

### RT-qPCR

RNA was extracted from aliquots of control and IFNγ stimulated splenocytes using RNeasy® Plus Mini Kit (Qiagen, Hilden, Germany, Cat #74134) following manufacturers’ instructions. cDNA was synthetized from 1µg of purified RNA using QuantiTect® reverse transcription kit (Qiagen, Toronto, Ontario, Canada). Quantitative RT-PCR reactions were carried out in CFX Connect Real-Time PCR Detection System (Bio-Rad, Canada) using SYBR Green Supermix (Bio-Rad, Mississauga, Ontario, Canada). The expression of indicated genes was measured using primers listed in Table S2. Each primer for qRT-PCR was validated by analyzing its standard curve and melt curve. Data was normalized to the expression of a housekeeping gene β-actin in unstimulated cells from *Nlrc5^+/+^* mice to calculate the relative gene-expression.

### Induction of Fibrosarcoma

To test the role of NLRC5 in cancer immune surveillance, endogenous fibrosarcoma was induced by subcutaneous administration of 3-methylcholanthrene (MCA; Sigma-Aldrich, Cat # 212942) in the flanks of *Nlrc5^-/-^*, *Nlrc5^+/+^* and *Rag1^-/-^*mice following published methods [24,52]. As the effective MCA dose varies among mouse colonies even for the same genetic strain [52,53], 100, 200 and 400 μg of MCA in 100 μL corn oil. Both male and female mice were used, as no significant difference was observed in tumor incidence between them. Tumor development was initially monitored visually once week and, after palpable tumor development, by using digital vernier calipers until the endpoint of 20 mm diameter in any direction or tumor skin ulceration, at which point the mice were euthanized. Tumor incidence, growth and endpoints were recorded for statistical analyses. Tumor tissues were collected, and tissue pieces were fixed in buffered formalin for paraffin embedding, snap frozen in liquid nitrogen for proteomic analysis and cultured to establish cancer cell lines.

### Histology and immunohistochemistry

Formalin fixed paraffin embedded (FFPE) tumor sections were deparaffinized, rehydrated, and stained with hematoxylin and eosin (H&E). For immunohistochemical detection of CD3, rehydrated tumor sections were immersed in citrate buffer (pH 6.0) and given intermittent microwave treatment to retrieve antigenic epitopes. Following incubation in 3% hydrogen peroxide for 10 min to inhibit endogenous peroxidase activity, sections were blocked with 5% BSA in Tris-buffered saline (TBS) containing 20% Tween-20 (TBS-T). Slides were incubated overnight at 4°C with a rat monoclonal antibody against mouse CD3 (ThermoFisher scientific, Cat #14-0032-81) diluted in blocking buffer, washed and then incubated with horseradish peroxidase (HRP)-conjugated secondary Ab for 1 h. After thorough washing, slides were incubated in 30-40 µL of signal stain boost (SignalStain^®^ Boost, Cell Signaling Technology, Cat # 8125) for 30 minutes at room temperature. After washing. a substrate solution containing 3,3’-diaminobenzidine (DAB; Sigma-Aldrich; 30 μL chromogen diluted in 1 mL of DAB liquid buffer) was added for 10 min. The sections were counterstained with hematoxylin and mounted with a coverslip. Images of the stained sections, digitized using the NanoZoomer Slide Scanner (Hamamatsu Photonics, Japan), were analyzed by the NanoZoomer Digital Pathology software NDPview2.0. Necrotic areas in H&E-stained sections were quantified using the NIH ImageJ software (version 1.53e) and CD3 positive cells were counted in NanoZoomer images.

### Assay for cancer immunoediting

Tumor tissues collected aseptically from individual tumors were minced in sterile PBS, digested with 1 mg/mL collagenase (type II, Cat #LS004176, Worthington, NJ, USA) and 40 μg/mL DNase I (Roche, Cat #10104159001) at 37°C for 60 mins. Clumps were removed by filtering through 70 µm membrane filter and cells were by centrifugation for 5 mins at 400*g*. Cells were resuspended in RPMI-1640 cell culture medium containing 10% fetal bovine serum, plated in adherent culture plates and passed through several passages over three months prior to using them for immunoediting experiments or freezing in liquid nitrogen. Each independent tumor derived cell line established from *Nlrc5^−/−^*, *Nlrc5^+/+^* and *Rag1^−/−^*mice was injected simultaneously into batches of *Rag1^−/−^* and C57BL/6 hosts. The cells (2 x 10^5^ cells in 50 μL PBS) were injected subcutaneously in the flanks. Tumor growth was monitored until the end point.

### Statistical Analysis

Data analysis and graphic plotting were carried out using the GraphPad Prism (San Diego, CA, USA; version 10.4.1). Log-rank test was used to calculate survival probability. For other comparisons between groups, one-way ANOVA was used. *p* values are represented by asterisks: * <0.05, ** <0.01, *** <0.001, **** <0.001.

### Mass spectrometry

#### Protein preparation and protease digestion

Snap frozen tumor tissues (20-50 mg) were resuspended in 1 mL of lysis buffer (8 M Urea, 1 M NH_4_HCO_3_ and 10 mM HEPES-KOH pH7.5) in a 2 mL low protein binding tubes (Axygen). Tissues were homogenized using steel beads in a mixer mill tissue lyser (MM 400, Retsch, Haan, Germany). Tissue lysates were transferred to fresh tubes and sonicated on ice (12 cycles, 20-25% intensity, 5 sec PULSE/ 5 sec OFF) and centrifuged at 16,000 *g* for 10 min at 4°C. Supernatants were transferred to fresh and proteins quantified using DC Protein Assay Kit (Bio-Rad, #5000113, #5000114, #5000115) following the manufacturer’s instructions. One hundred µg of protein was transferred to new tubes, volumes adjusted to 50 µL with urea solution (8 M Urea, 10 mM HEPES, pH 8) and 1 µL of 255 mM dithiothreitol was added to achieve 5 mM final concentration. The tubes were vortexed and boiled at 95°C for 2 min and allowed to cool at room temperature for 30 min. After adding 1.5 µL of photosensitive 262.5 mM chloroacetamide to achieve 7.5 mM final concentration, the samples were vortexed and incubated in dark at room temperature for 20 min before adding 150 µL of 50 mM NH_4_HCO_3_ to bring the urea concentration down to 2 M. Proteins were digested by adding 1 µg of Pierce^TM^ Trypsin Protease (Thermo Scientific, cat# 90058) and overnight incubation at 30°C. Proteolysis was stopped by adding 0.5 µL of 100% Trifluoroacetic acid (TFA) and the digested peptides were cleaned using Pierce^TM^ C18 tips (Thermo Scientific, cat# 87784). The solvents were then removed using Vacufuge Plus centrifuge concentrator (Eppendorf) and the peptides diluted in 1% Formic Acid (FA) were quantified using NanoDrop 2000/2000c spectrophotometer (Thermo Fisher scientific).

#### Liquid chromatography-tandem mass spectrometry (LC-MS/MS)

Concentrated peptides (250 ng) were separated on a nanoHPLC system (nanoElute, Bruker Daltonics). The samples were loaded onto an Acclaim PepMap100 C18 Trap Column (0.3 mm id x 5 mm, Dionex Corporation) at 4 µL/min consistent flow and peptides were eluted onto a PepMap C18 analytical nanocolumn (1.9 µm beads size, 75 µm x 25 cm, PepSep) heated at 50°C. Peptides were eluted with solvent B (100% ACN & 0.1% FA) in a 5-37% linear gradient with a flowrate of 400 nL/min for ∼2 h. The HPLC system was coupled to an TimsTOF Pro ion mobility mass spectrometer containing Captive Spray nano electrospray source (Bruker Daltonics). Data acquisition was done using diaPASEF mode. For each individual Trapped Ion Mobility Spectrometry (TIMS) measurement in diaPASEF mode, a single mobility window consisting of 27 mass steps (with m/z ranging from 114 to 1414 and a mass width of 50 Da) was employed per cycle, which had a 1.27-second duty cycle. This process involves scanning the diagonal line in the m/z-ion mobility plane for +2 and +3 charged peptides.

#### Protein identification

Peptide mass spectra were analyzed using the DIA-NN, an open-source software [54] suite for DIA / SWATH data processing (https://github.com/vdemichev/DiaNN, version 1.8.1), installed in an Apptainer container (https://apptainer.org/, version 1.3.5) using docker image provided on the docker hub [55]. Analysis was performed using default parameters except for these options: 2 missed - cleavages was allowed; trypsin digestion was performed for K/R; protein N-term methionine excision as variable modification for the *in-silico* digest.

The *Mus musculus* reference proteome UP000000589 was downloaded from the Uniprot website (https://www.uniprot.org/proteomes/UP000000589). The reference proteome contained a total of 63367 proteins. For the FASTA search, DIA-NN was instructed to perform an *in-silico* digest of the sequence database. A mass tolerance accuracy of MS1 and MS2 of 20 ppm was used for precursor and fragment ions, respectively. Minimum and maximum were set for peptide length (7-30 amino acids), and precursor charge (1-5), precursor m/z (100-1700) and fragmentation m/z (100-1500) for *in silico* library generation or library-free search. For the reanalysis, MBR (match between run) was enabled and chosen the smart profiling when creating a spectral library from DIA data. Carboxyamidomethylation (unimod4), and oxidation (M) (unimod35) were set as fixed modifications and N-terminal protein acetylation was set as a variable modification. DIANN protein group matrix was filtered using custom Perl script to extract protein groups with a single protein. This new matrix was used as input to the R package FragPipeAnalystR [56] to get QC metrics and differential protein expression profiles.

#### Proteomic data visualization

Significantly modulated proteins with cut-off values of log2-fold change <−1 and >1 and *p*-Value <0.05 in the FragPipeAnalystR pipeline output were sorted using Microsoft Excel (Office 365). GraphPad Prism version 10.0.3 (GraphPad, Boston, MA) was used to generate volcano plots and pie charts. Venn diagrams were generated using the jvenn https://jvenn.toulouse.inrae.fr/app/index.html; accessed in Dec 2024) online tool [57]. The SRplot server (http://www.bioinformatics.com.cn/srplot; accessed in Dec 2024) was used for pathway and Gene Ontology (GO) analyses and to generate pathway enrichment plots and heatmaps. Gene set enrichment analysis (GSEA) based on GO and Kyoto Encyclopedia of Genes and Genomes (KEGG) was conducted using HTSanalyzeR2 (https://github.com/CityUHK-CompBio/HTSanalyzeR2) [58].

## Results

### Increased incidence of MCA-induced fibrosarcoma and reduced survival in Nlrc5^−/−^ mice

To elucidate the role of NLRC5 in tumor immunosurveillance, we injected 3-methylcholanthrene (MCA) into *Nlrc5^−/−^* mice, using *Nlrc5^+/+^* and *Rag1^−/−^* mice as controls. All mice are in C57BL/6 background. Different doses of MCA (100μg, 200μg, and 400μg) were used because susceptibility of the same genetic strain to MCA-induced tumors can vary from one animal facility to another [52]. At 100μg MCA, 7 out of 8 *Nlrc5^−/−^* mice developed tumors compared to six out of nine *Nlrc5^+/+^* mice, with *Nlrc5^−/−^* mice with higher tumor-related mortality (87.5% versus 70.5%) (Figure 1A, B). Seven out of eight *Nlrc5^−/−^*mice developed tumors, compared to four out of nine *Nlrc5^+/+^* mice at 200μg MCA, again with significantly higher mortality than in *Nlrc5^+/+^*mice (87.5% versus 55.5%) (Figure 1C, D). At 400μg MCA, all *Nlrc5^−/−^* mice and three out of five *Nlrc5^+/+^* mice developed tumors (mortality 100% versus 60%) (Figure 1E, F). *Rag1^−/−^*mice developed tumors at all doses of MCA tested. Pooled data from all three doses of MCA showed that 91.3% of *Nlrc5^−/−^* mice developed tumors compared to the 56.5% observed in *Nlrc5^+/+^*mice, with significantly elevated mortality (91.3% versus 69%) (Figure 1G, H). These findings indicate that NLRC5 expression is required for efficient control of endogenously arising tumors and underscore the key role of NLRC5 in cancer immunosurveillance.

**Figure 1.**
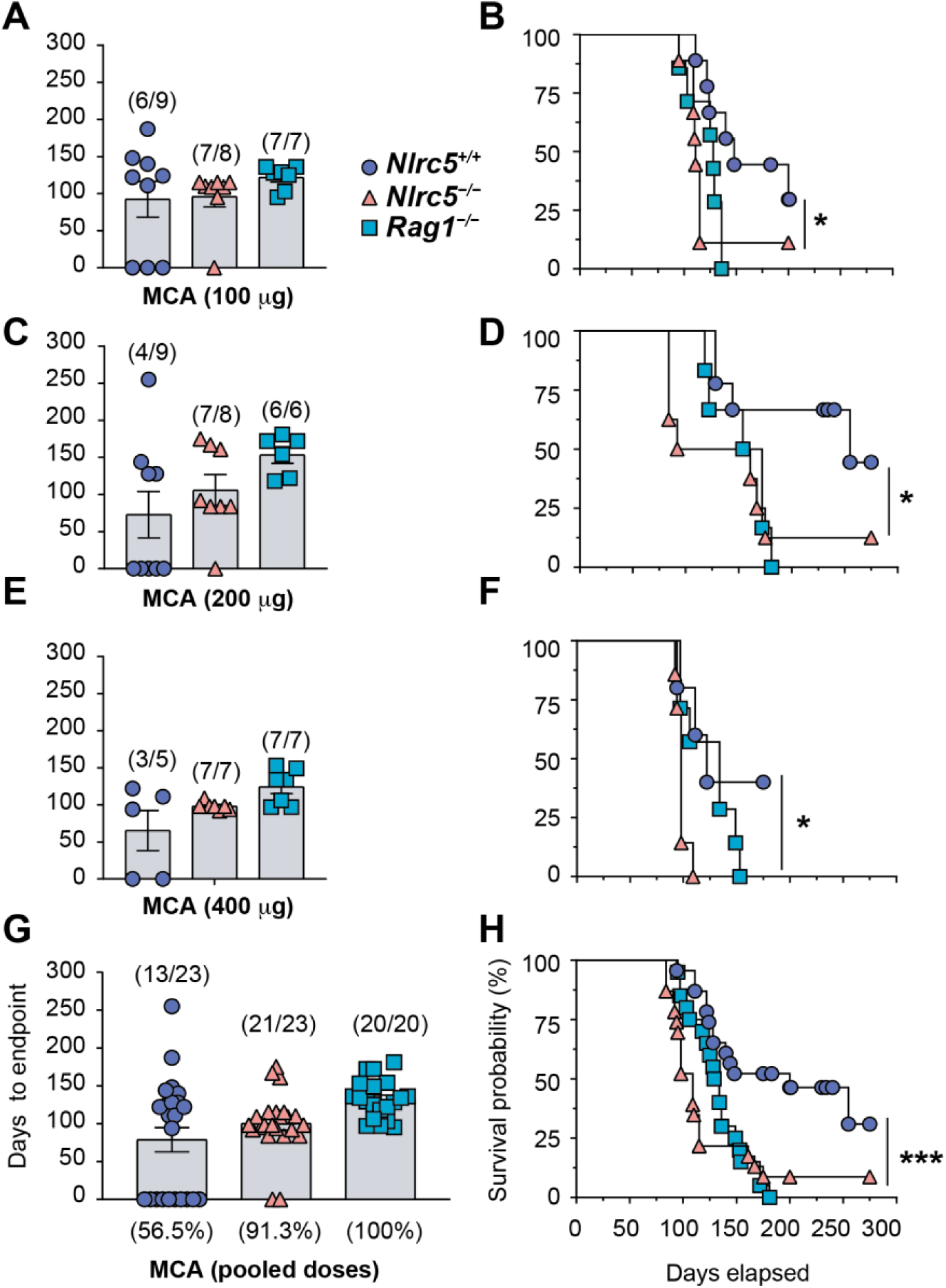
NLRC5 deficiency increases susceptibility to MCA-induced fibrosarcoma. *Nlrc5^−/−^*, *Nlrc5^+/+^* and *Rag1^−/−^* mice were injected the chemical carcinogen 3-methylcholanthrene (3-MCA) at different doses of 100 μg (A, B), 200 μg (C, D), 400 μg (E, F) subcutaneously. Development of fibrosarcoma was monitored visually and by using a digital vernier caliper until the endpoint (tumor diameter of 20 mm in any direction, or ulceration at any diameter), when the animals were euthanized. Days to endpoint are plotted in the left panels (A, C, E) and survival probability in the right panels (B, D, F). Pooled data are shown in G and H. Numbers in parenthesis in A, C, E and G indicate the number of mice that developed tumors over total number for each genotype. B, D, F, H: Log-rank (Mantel-Cox) test. * *p* ≤0.05, *** *p* ≤0.001.

### Histological features of MCA-induced fibrosarcoma in Nlrc5^−/−^ and control mice

Histological examination of tumor tissues revealed the distinctive feature of fibrosarcoma that is characterized by spindle-shaped fibroblast-like cells arranged in a herringbone or chevron-like pattern (Figure 2A). Notably, tumors arising in *Nlrc5^+/+^*and *Nlrc5^−/−^* mice displayed large areas of tumor necrosis that did not occur in *Rag1^−/−^* tumors (Figure 2A, B), suggesting cell death mediated by the immune system rather than spontaneous death caused, for example, by hypoxia or reduced nutrient supply. This notion is supported by significant infiltration by CD3+ T lymphocytes in tumors from *Nlrc5^+/+^* and *Nlrc5^−/−^* mice but not in *Rag1^−/−^* hosts (Figure 2C, D).

**Figure 2.**
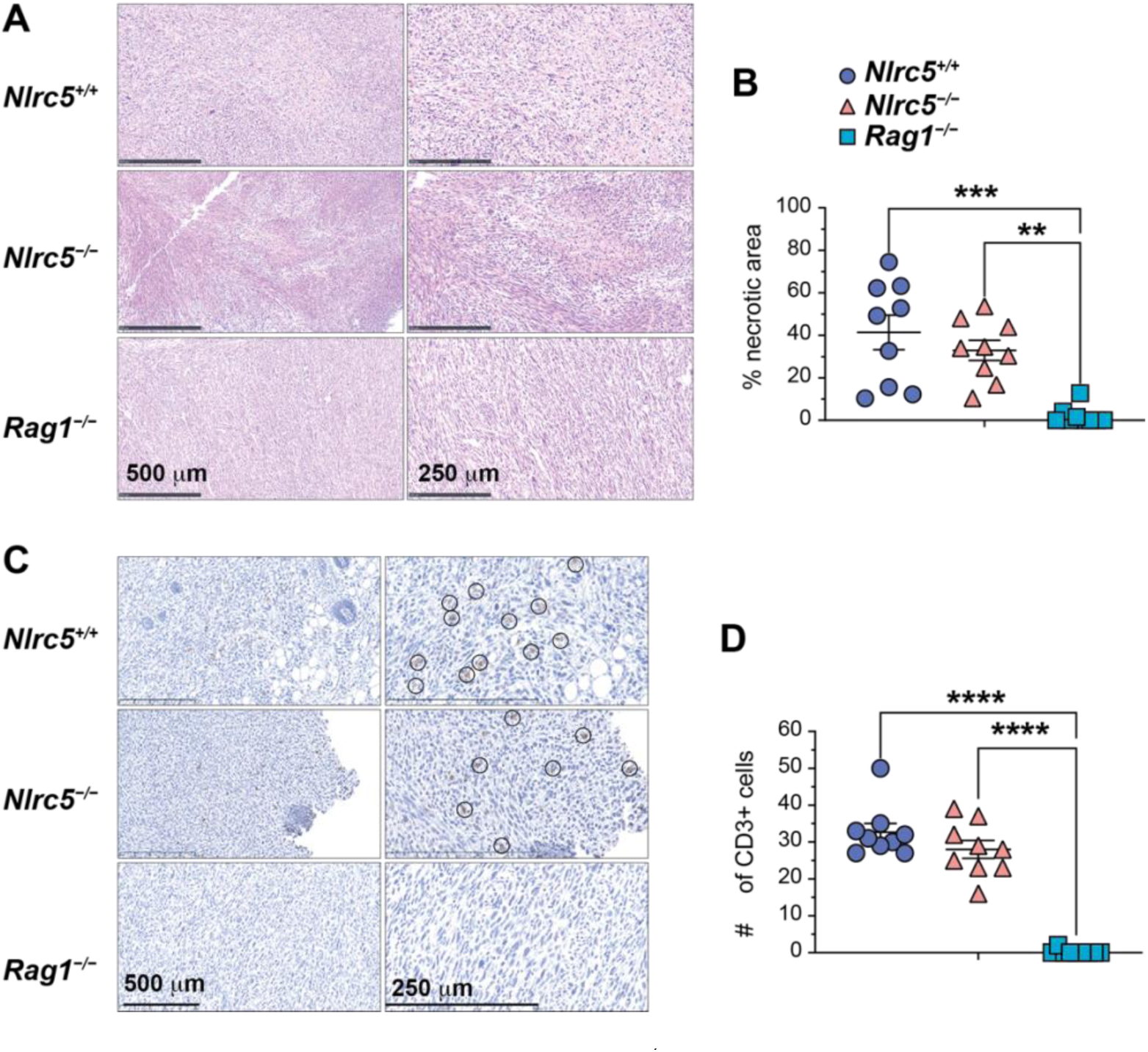
MCA-induced tumors in *Nlrc5^−/−^* mice show areas of necrosis and T cell infiltration as in *Nlrc5^+/+^* mice. (A) H&E staining of MCA-induced tumors from *Nlrc5^+/+^*, *Nlrc5^−/−^* and *Rag1^−/−^*mice at lower and higher magnification, the latter revealing the Herringbone pattern. Tumors from *Nlrc5^+/+^* and *Nlrc5^−/−^*mice display pale staining areas of necrosis that are not observed in *Rag1^−/−^*tumors. Representative data from three mice in each genotype. (B) The proportion of necrotic area in random fields of three tumors per genotype was measured using ImageJ. (C) Immunohistochemical staining of MCA-induced tumors for CD3^+^ T cells (brown-stained cells indicated by circles). Representative data from 3 mice in each group. (D) The number of CD3^+^ T cells were counted and pooled from 3 random fields per tumor. Mean + SEM; One-way ANOVA ** *p* ≤0.05, *** *p* ≤0.001, **** *p* ≤0.0001.

### NLRC5 is required for cancer immunoediting

To investigate the immunoedited status of MCA tumors, cancer cell lines were established from the MCA-induced tumors (Figure 1). Independent cell lines from *Nlrc5^+/+^*, *Nlrc5^−/−^* and *Rag1^−/−^* tumors were implanted into immunocompetent C57BL/6 and immunocompromised *Rag1^−/−^* mice, and their growth was assessed. Cell lines from *Rag1^−/−^* tumors serve as controls for non-immunoedited status, whereas cell lines from *Nlrc5^+/+^* tumors represented immunoedited status as shown by Shankaran et al., [24]. As expected, the two *Rag1^−/−^* tumor cell lines showed unhindered growth when implanted into a new batch of *Rag1^−/−^* hosts (Figure 3A). However, when implanted into C57BL/6 hosts, the same tumor cell lines were efficiently controlled because they had not previously been controlled by the adaptive immune system, i.e., not immunoedited (Figure 3A). On the other hand, five distinct MCA cancer cell lines originating from the MCA-induced tumors of *Nlrc5^+/+^* hosts showed unhindered growth in fresh cohorts of C57BL/6 mice (25 out of 26 mice; only three cell lines are shown in Figure 3B), indicating their immunoedited status that was previously established in *Nlrc5^+/+^*hosts. Notably, for reasons that remain unclear, *Nlrc5^+/+^* tumor cell lines frequently ulcerated in *Rag1^−/−^* hosts, contributing to their earlier endpoints (Figure 3B).

**Figure 3.**
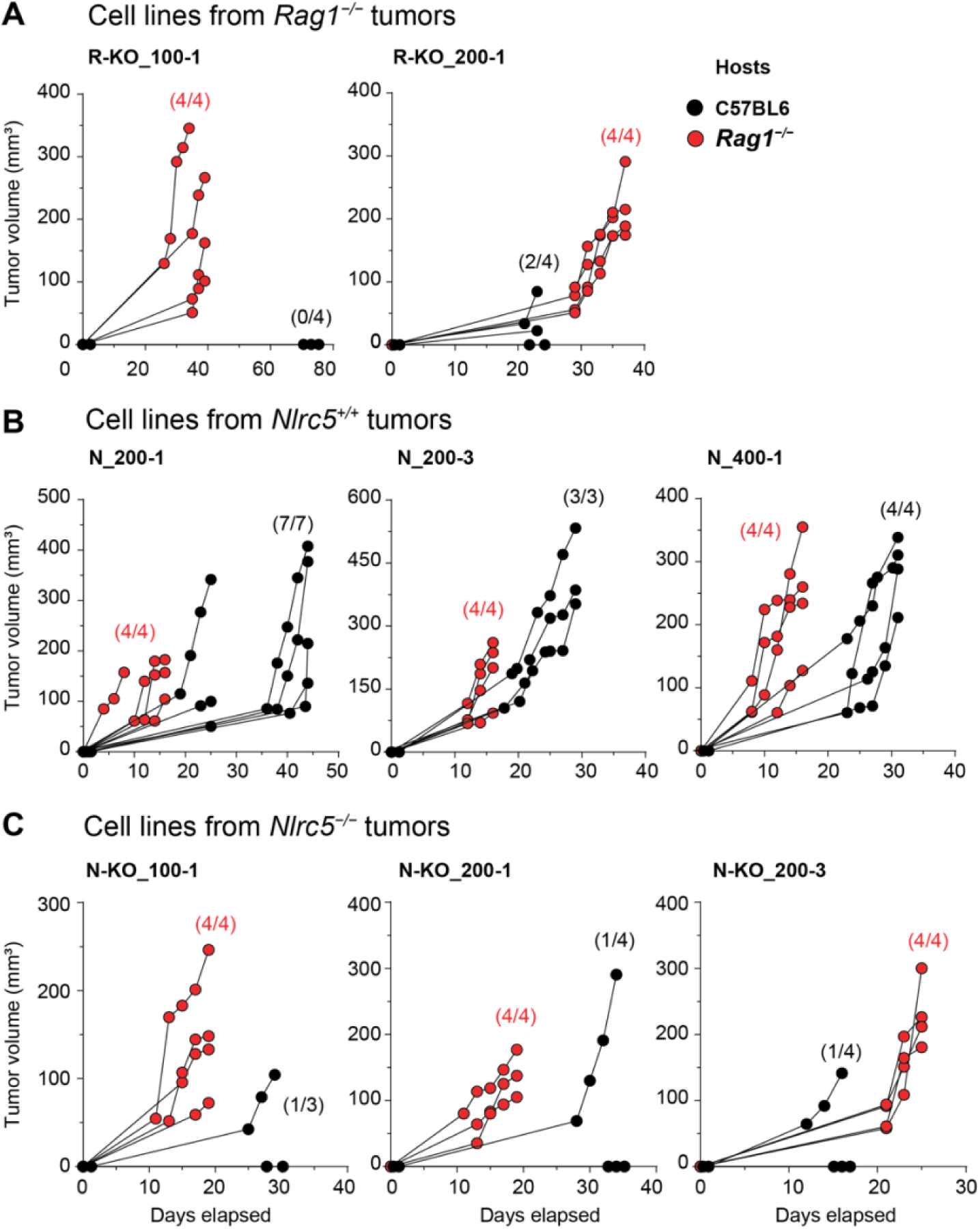
Impaired immunoediting of MCA tumors in *Nlrc5^−/−^*mice. Cancer cell lines established from MCA-induced fibrosarcoma were implanted into immunocompetent C57BL/6 (black circles) and immunocompromised *Rag1^−/−^*(red circles) hosts (200,000 cells in 50 µl 1xPBS, subcutaneous route). Tumor growth was monitored every 2 days. The final data points indicate the endpoint. (A) Tumors developing in *Rag1^−/−^* mice were not immunoedited. *Rag1^−/−^* tumor cell lines arising from 100 or 200 µg MCA are indicated as R-KO_100-1 and R-KO_200-1. (B) Tumors developing in *Nlrc5^+/+^* hosts were immunoedited. Three of the five cell lines established from MCA-induced fibrosarcoma of *Nlrc5^+/+^* hosts injected with 100 or 200 µg MCA are indicated as N_200-1, N_200-3 and N_400-1. (C) Impaired immunoediting of MCA tumors in *Nlrc5^−/−^* mice. Three of the four *Nlrc5^−/−^* tumor cell lines arising from 100 or 200 µg MCA-induced tumors are indicated as N-KO_100-1, N-KO_200-1 and N-KO_200-3.

Evaluation of the immunoedited status in MCA tumors arising in *Nlrc5^−/−^* mice using four different cancer cell lines showed that the majority of *Nlrc5^−/−^*tumor cell lines were effectively controlled in C57BL/6 hosts with only a small proportion of mice developing tumors (5 out of 15) (only three cell lines are shown in Figure 3C). Most of these cell lines grew uninhibited in *Rag1^−/−^* hosts, with all 15 mice developing tumors. These data show that unlike tumors arising in *Nlrc5^+/+^* mice that are immunoedited, most tumors arising in *Nlrc5^−/−^* mice were non-immunoedited. These findings also suggest that the NLRC5-dependent tumor growth mechanisms are the key contributors to cancer immunoediting.

### Impact of NLRC5 deficiency on the tumor proteome

NLRC5 deficiency compromised tumor growth control and tumor immunoediting to a level comparable to *Rag1^−/−^* mice (Figures 1, 3), indicating that NLRC5-dependent adaptive antitumor immune mechanisms are critical for tumor growth control. To gain deeper understanding of these control mechanisms, we carried out mass spectrometry analysis of the proteomes from three independent MCA tumors for each of the three genotypes. A total of 6373 individual proteins were identified in these samples, and quality control measures showed a random missing protein profile (Figure 4A, B). Even though Pearson’s correlation coefficient revealed high degree of similarity among the nine tumors, presumably because they all represent fibrosarcoma (Figure 4C). PCA plot showed that the tumors did not tightly cluster according to the three genotypes (Figure 4D), possibly reflecting the randomness of the oncogenic pathways activated in these tumors by the chemical carcinogen MCA.

**Figure 4.**
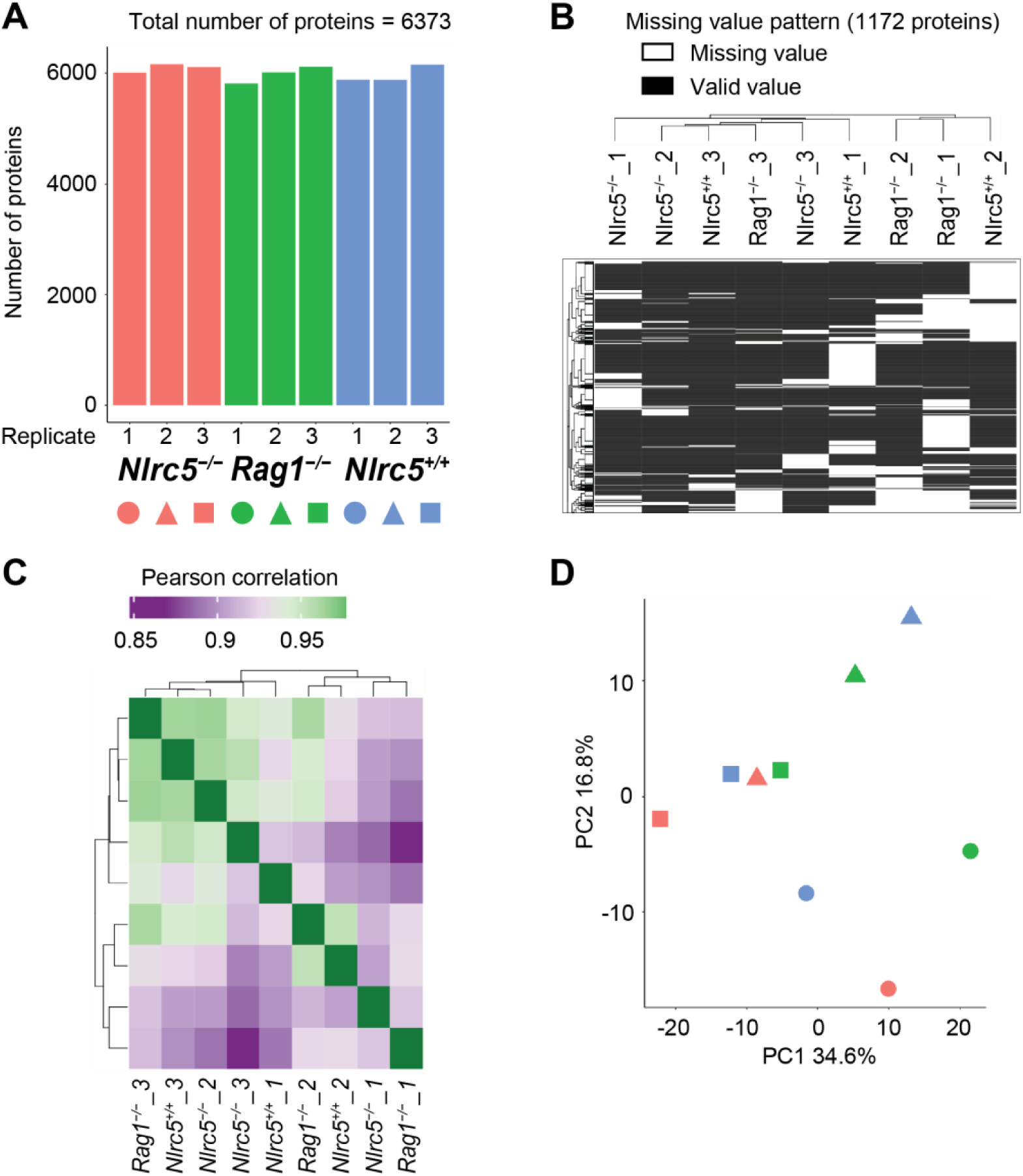
Quality control for mass spectrometry data on tumor proteomes. Three independent MCA tumors (biological triplicates) from each of the *Nlrc5^+/+^*, *Nlrc5^−/−^*and *Rag1^−/−^* mice were subjected to shotgun proteomics by LC-MS/MS. (A) Number of individual proteins identified in each tumor by more than peptide. (B) Missing value pattern across samples. (C) Pearson correlation between samples. (D) Principal component analysis (PCA) plot.

Pairwise comparison of protein expression profiles using Volcano plots identified only 239 differentially expressed proteins (DEPs) between tumors arising in *Nlrc5^+/+^* mice and *Nlrc5^−/−^* mice (out of 6373 proteins), with 140 upregulated proteins (green) and 99 downregulated proteins (red) in *Nlrc5^+/+^*tumors (Figure 5A). Similar comparison between *Nlrc5^+/+^* and *Rag1^−/−^* tumors revealed 171 DEPs with 82 upregulated and 89 downregulated proteins in *Nlrc5^+/+^* tumors (Figure 5B). *Nlrc5^−/−^* and *Rag1^−/−^* tumors showed an elevated number of DEPs (357), with 116 upregulated and 241 downregulated proteins in *Nlrc5^−/−^* tumors (Figure 5C), suggesting that *Nlrc5^−/−^* tumors are more disparate from *Rag1^−/−^* tumors than from *Nlrc5^+/+^* tumors. To further understand these differences, the degree of overlap between differentially modulated proteins was analyzed using a Venn diagram (Figure 5D). The three groups of DEPs shared only 6 proteins (CPS1, RABEPK, ARHGEF5, TNC, TPM1, AND MUP3), indicating that fibrosarcoma arising in *Nlrc5^+/+^*, *Nlrc5^−/−^* and *Rag1^−/−^* mice display a subset of distinct proteins that may be associated with genotype-specific host response toward the tumor rather than with inter-tumor heterogeneity. Among the 239 DEPs between *Nlrc5^+/+^* versus *Nlrc5^−/−^* tumors, only 33 were shared with the DEPs of *Nlrc5^+/+^* versus *Rag1^−/−^*tumors (Figure 5D), suggesting that loss of NLRC5 or RAG1 differentially impact the MCA tumor proteome. This notion is further supported by 81 out of 357 DEPs between *Nlrc5^−/−^* versus *Rag1^−/−^* tumors being shared with the DEPs of *Nlrc5^+/+^* versus *Nlrc5^−/−^* tumors, compared to 45 shared with the DEPs of *Nlrc5^+/+^* versus *Rag1^−/−^*tumors (Figure 5D). Notably, 225 DEPs between *Nlrc5^−/−^* versus *Rag1^−/−^* tumors were unique compared to 119 unique DEPs for *Nlrc5^+/+^* versus *Nlrc5^−/−^* tumors and 83 unique DEPs for *Nlrc5^−/−^* versus *Rag1^−/−^*tumors, suggesting that NLRC5 deficiency differentially impacts the host response to endogenously arising tumors.

**Figure 5.**
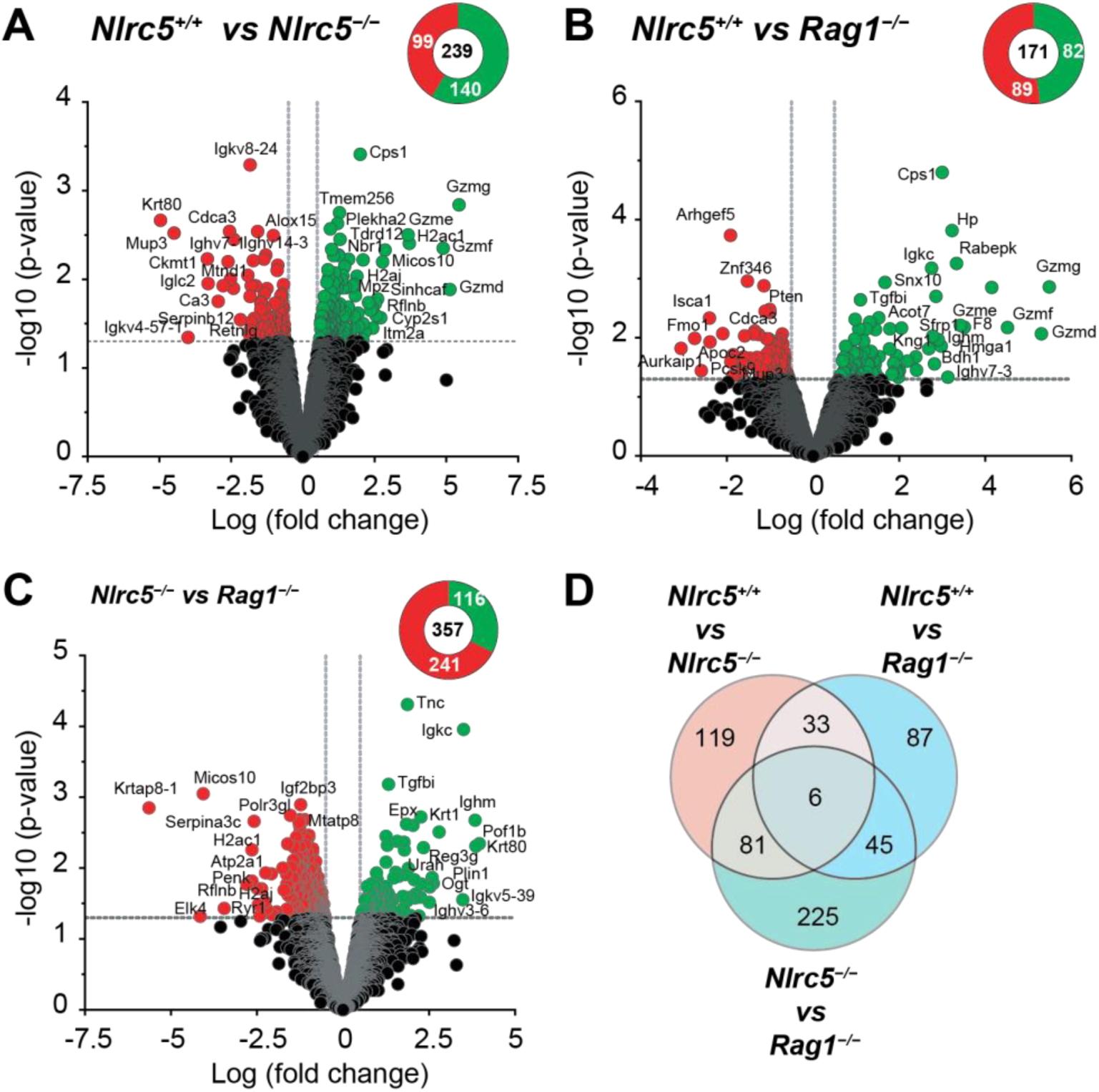
Differential protein expression in MCA tumors. (A-C) Volcano plots showing significantly modulated proteins between *Nlrc5^+/+^*and *Nlrc5^−/−^* tumors (A), *Nlrc5^+/+^* and *Rag1^−/−^*tumors (B) *Nlrc5^−/−^* and *Rag1^−/−^* tumors (C); vs, versus. Differentially expressed proteins (DEPs; fold change >1.5 fold; *p* <0.05) are indicated in green (upregulated) or red (downregulated). Total number of significantly upregulated and downregulated proteins are shown in the pie chart. (D) Venn diagram showing unique and shared DEPs modulated by the loss of NLRC5 or RAG1 in the host.

### Enrichment of CTL-mediated cell death pathways in Nlrc5^+/+^ tumors

To elucidate the functional significance of proteins modulated by NLRC5 deficiency in MCA tumors, we carried out Gene ontology (GO) analysis of DEPs for potential impact on biological processes (BP), cellular compartments (CC), and molecular functions (MF) (Figure 6). DEPs upregulated in *Nlrc5^+/+^* tumors in comparison to *Rag1^−/−^*tumors showed enrichment of BP terms such as granzyme-mediated apoptotic signaling pathway, positive regulation of inflammatory response and antigen processing and presentation, as well as the MF terms CD8 receptor binding and T cell antigen receptor binding (Figure 6A). The BP term granzyme-mediated apoptotic signaling pathway and the MF term NF-κB binding were enriched among the DEPs upregulated in *Nlrc5^+/+^* tumors in comparison to *Nlrc5^-/-^* tumors (Figure 6B), in line with the known functions of NLRC5 in MHC-I-dependent CD8 T cell activation and regulation of NF-κB activity [31,59]. None of these terms were represented among those enriched within the upregulated DEPs of *Nlrc5^-/-^* tumors in comparison to *Rag1^−/−^* tumors (Figure 6C). Heatmap plot of proteins involved in ‘granzyme-mediated apoptotic signaling pathway’, ‘granzyme-mediated programmed cell death signaling pathway’ and ‘cytolysis’ showed upregulation of several murine granzymes (D, E, F, G) that play a crucial role in CTL-mediated cytotoxicity [60] in *Nlrc5^+/+^* tumors compared to *Nlrc5^−/−^*and *Rag1^−/−^* tumors (Figure 7A). Corroborating with this data, gene set enrichment analysis (GSEA) of DEPs showed that *Nlrc5^+/+^* tumors displayed high enrichment score (0.795) for proteins involved in CD8 receptor binding (GO: 0042610) that included classical (H-2D1, H2-K1) and non-classical (H2-Q4, H2-Q7) MHC-I molecules (Figure 7B, C). Notably, this group of CD8 receptor binding proteins included FYN, which is downregulated in *Nlrc5^+/+^* tumors compared to *Nlrc5^−/−^*and *Rag1^−/−^* tumors (Figure 7C). Collectively, these results indicate that CTL-mediated cytotoxic pathway is the key mediator of NLRC5-dependent control of endogenously arising tumors.

**Figure 6.**
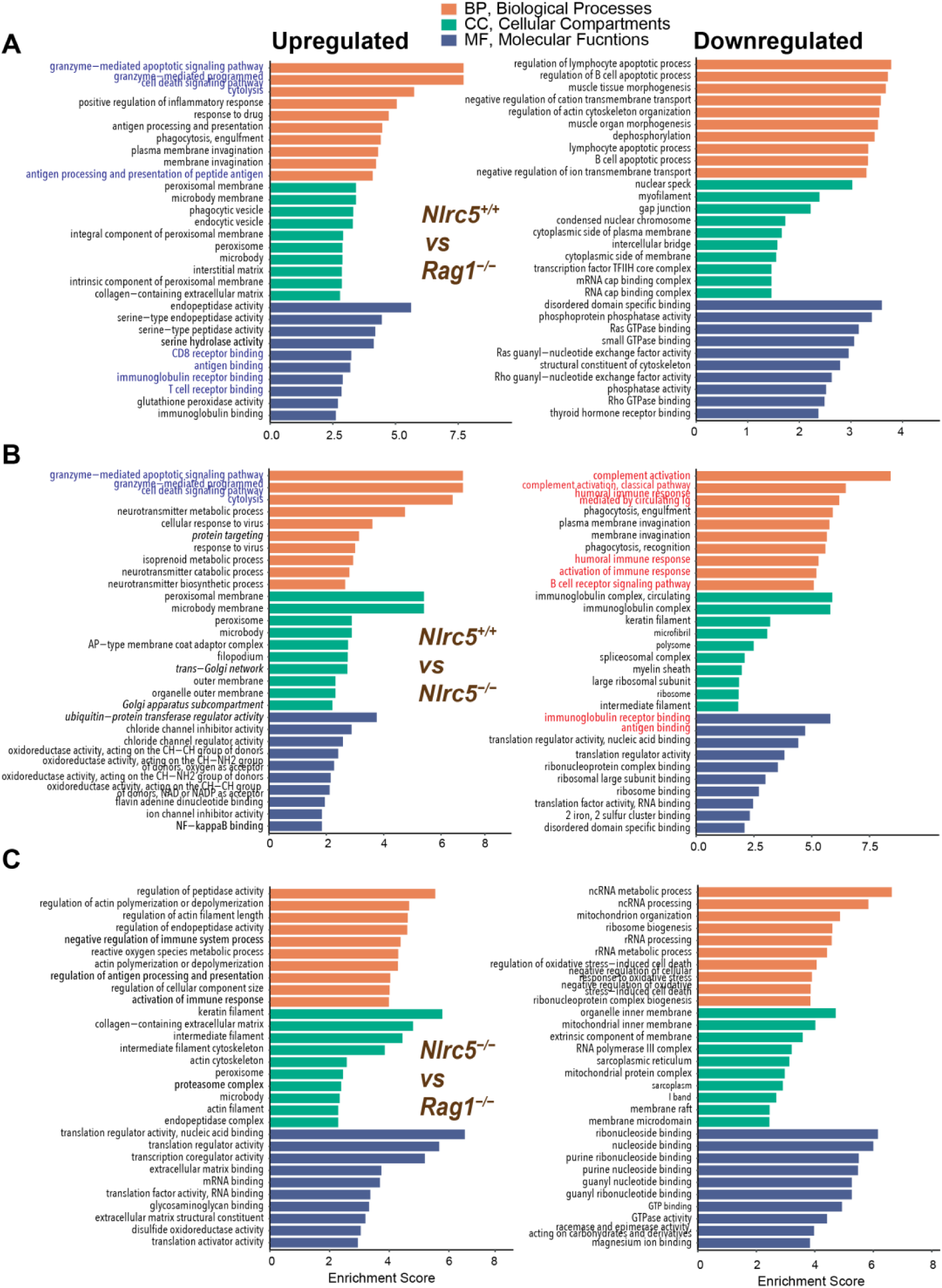
Gene ontology analysis of DEPs in MCA tumors. Gene ontology (GO) analysis of upregulated and downregulated genes in *Nlrc5^+/+^* versus *Rag1^−/−^*tumors (A), *Nlrc5^+/+^* versus *Nlrc5^−/−^* tumors (B), and *Nlrc5^−/−^* versus *Rag1^−/−^* tumors (C); vs, versus. GO terms in Biological Processes (BP, orange bars), Cellular Compartments (CC, green bars), and Molecular Functions (MF, blue bars) are indicated with corresponding enrichment scores. GO terms related to tumor control by cytolytic immune cell function are indicated in blue color fonts, and terms related to innate immune response in red color font.

**Figure 7.**
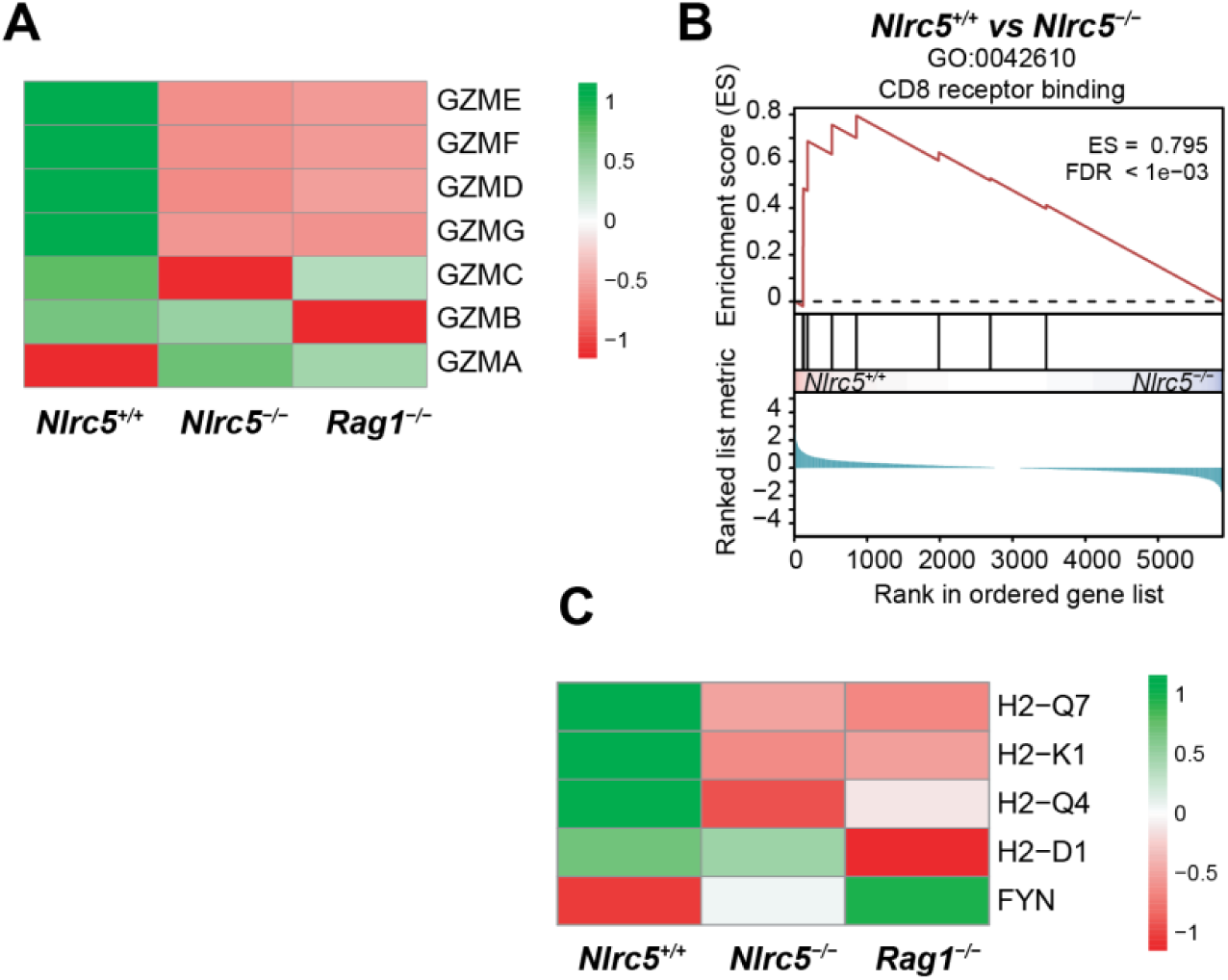
Expression level of proteins identified within the key ontology terms upregulated in *Nlrc5^+/+^* tumors. (A) Heatmap analysis shows the expression level of proteins within the BP terms granzyme mediated elimination and cytolysis. (B, C) GSEA plot (B) and corresponding heatmap (C) for proteins identified within the MF term CD8 receptor binding. (A, C) Normalised abundance values (calculated using complete-linkage clustering and Euclidean distance method) are color coded on a scale with green and red representing upregulation and downregulation, respectively.

### Upregulation of innate immune pathways in NLRC5 deficient tumors

Even though cytolytic immune response pathways were not enriched in *Nlrc5^−/−^* tumors, they displayed areas of necrosis that are not found in *Rag1^−/−^* tumors, suggesting activation of other cytolytic immune cell functions. Among the DEPs downregulated in *Nlrc5^+/+^*tumors in comparison with *Nlrc5^-/-^* tumors, but not in *Rag1^−/−^* tumors (Figure 6C), humoral immune response and innate immune response pathways were notably enriched. GSEA of DEPs in *Nlrc5^−/−^* versus *Rag1^−/−^* tumors revealed enrichment in immune-related pathways, particularly innate immune response in *Nlrc5^−/−^* tumors (GO: 0006695, 0002376, 0045087; Figure 8A, B, C). A heatmap analysis of 118 genes associated with these immune response pathways confirmed that *Nlrc5^−/−^* tumors displayed elevated expression of innate immune response proteins that surpassed their expression levels in *Nlrc5^+/+^* tumors (Figure 8D). In fact, *Nlrc5^−/−^* tumors showed high enrichment of immunoglobulin-mediated immune response (GO:0016064) against both *Rag1^−/−^*tumors (enrichment score 0.752) and *Nlrc5^+/+^* tumors (enrichment score 0.709) and (Figure 9A). Heatmap analysis revealed that multiple immunoglobulin heavy chain variable region (IGHV)-derived peptides were upregulated in *Nlrc5^−/−^* tumors (Figure 9B), suggesting upregulation of antibody-mediated tumor resistance mechanisms to compensate for the impaired CTL-mediated immune surveillance functions caused by the loss of NLRC5 or RAG1.

**Figure 8.**
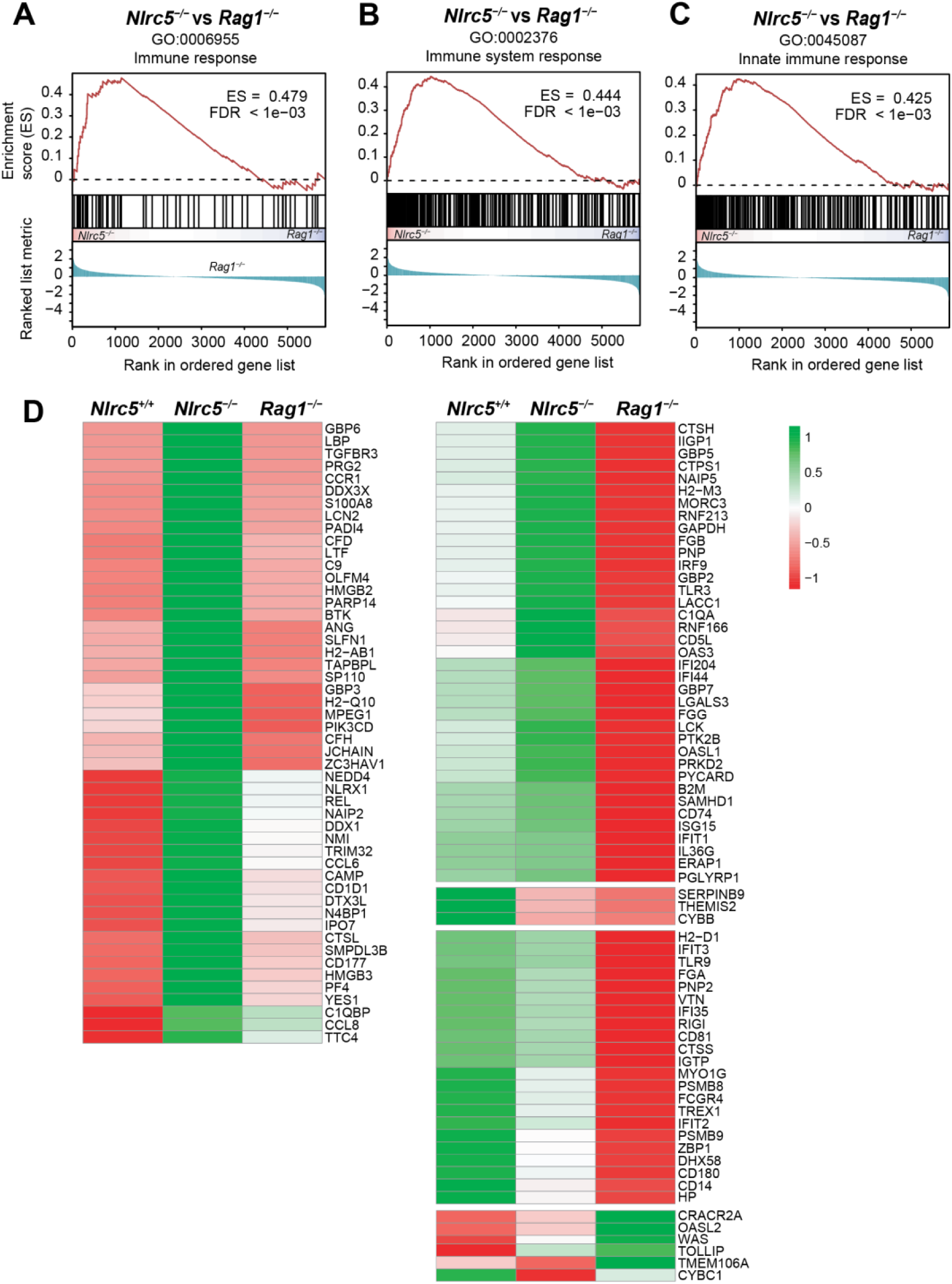
Upregulation of innate immune response pathways in *Nlrc5^−/−^* tumors: (A, B, C) GSEA of DEPs in *Nlrc5^−/−^*versus *Rag1^−/−^* tumors showing enrichment of immune response pathways. (D) Heatmap plot showing differential expression of proteins identified within these pathways in *Nlrc5^+/+^*, *Nlrc5^−/−^* and *Rag1^−/−^*tumors. Normalised abundance values (calculated using complete-linkage clustering and Euclidean distance method) are color coded on a scale with green and red representing upregulation and downregulation, respectively.

**Figure 9.**
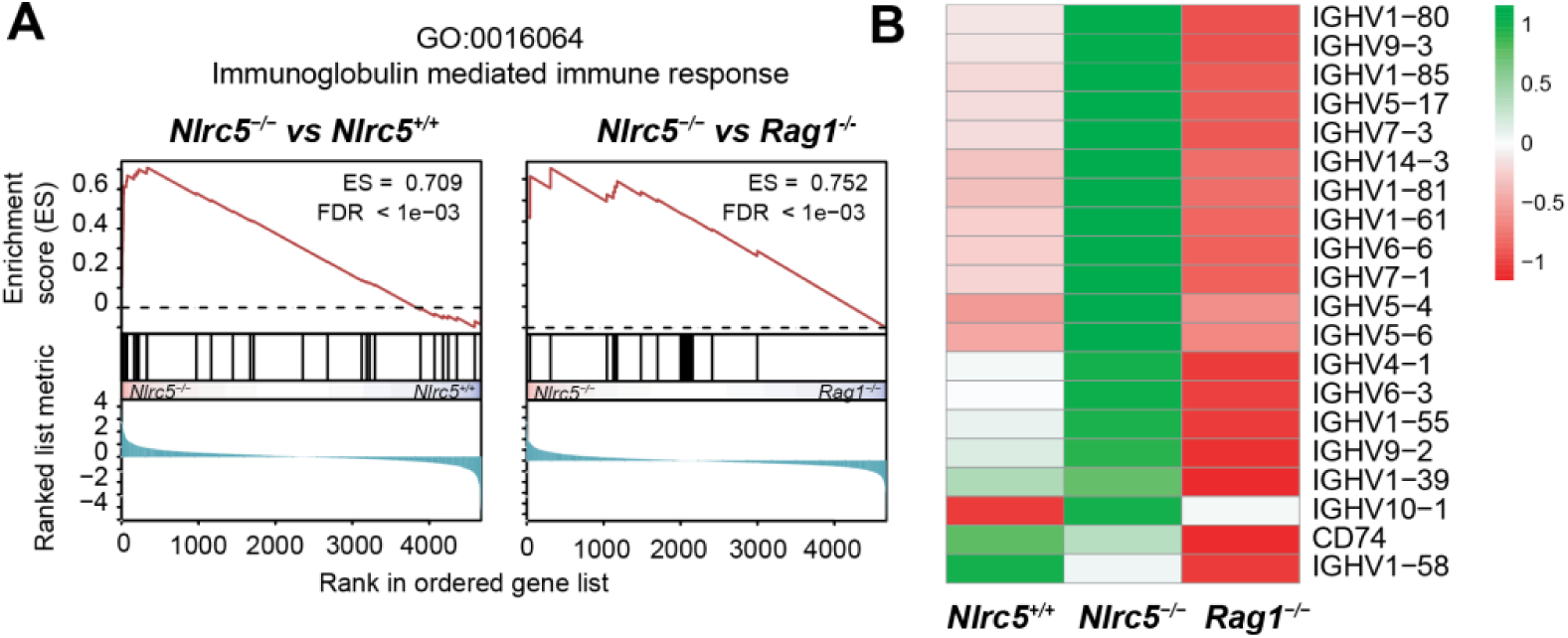
Enrichment of immunoglobulin-mediated immune responses in *Nlrc5^−/−^* tumors. (A) GSEA of DEPs in *Nlrc5^−/−^* tumors versus *Nlrc5^+/+^* or *Rag1^−/−^* tumors showing enrichment of proteins related to immunoglobulin-mediated immune response (GO:0016064). (B) Expression levels of proteins visualized using a heatmap. Normalised abundance values (calculated using complete-linkage clustering and Euclidean distance method) are color coded on a scale with green and red representing upregulation and downregulation, respectively.

DEPs upregulated in *Nlrc5^-/-^* tumors when compared to *Rag1^−/−^* tumors were also enriched for the GO terms regulation of peptidase activity, antigen processing and proteasome complex (Figure 6C), suggesting compensatory upregulation of pathways related to antigen processing and presentation. Furthermore, DEPs upregulated in *Nlrc5^-/-^* tumors in comparison to *Rag1^−/−^* tumors showed enrichment of the GO terms protein targeting (BP), trans-golgi network (CC) and ubiquitin-protease transferase activity (MF), suggesting that loss of NLRC5 may also trigger antigen presentation pathways outside of the proteasome machinery in an attempt to restore impaired CTL functions.

## Discussion

Using mouse genetic models, we have shown in this study that NLRC5 is required to reduce the incidence of endogenous tumors and to attenuate tumor progression. We have also shown that the tumors that arose in the absence of NLRC5 were not immunoedited similar to those arising in RAG1- deficient mice, underscoring the importance of NLRC5-dependent adaptive immune responses in delaying the progression of NLRC5 expressing tumors. Most of the NLRC5-dependent cancer immune surveillance and cancer immunoediting functions are likely mediated by the already demonstrated role of NLRC5 in transcriptional upregulation of MHC-I and key proteins involved in antigen processing and presentation to CD8^+^ T cells [33,48,49,61]. However, slightly faster tumor progression in *Nlrc5^-/-^* mice than in *Rag1^-/-^* mice, and notable differences in the proteomes of tumors arising in *Nlrc5^-/-^*mice compared to those of *Nlrc5^+/+^* and *Rag1^-/-^* tumors suggest that NLRC5 may also promote additional antitumor immune functions and possibly regulate potentially tumor-promoting immune responses.

A key step in the cancer immunity cycle [62,63], which involves activation of antitumor T lymphocytes and killing of tumor cells by CTLs in iterating cycles to achieve tumor growth control, is the presentation of tumor antigenic peptides to naïve CD8^+^ T cells. This process requires cross-presentation of tumor antigens acquired from dead tumor cells by dendritic cells (DC), which present the tumor antigenic peptides along with costimulatory ligands in tumor draining lymph nodes [64,65]. DCs can also acquire MHC-I:peptide complexes directly from live tumor cells via trogocytosis, and recent studies show that such cross-dressing of DCs with tumor cell-derived MHC-I:peptide complexes is important for initiating antitumor CTL responses [66–68]. DCs from *Nlrc5^-/-^* mice also show reduced MHC-I expression and are less efficient in activating CD8^+^ T cells [39,40,69]. As we have used in our studies whole body NLRC5 knockout mice, the aggressive growth of MCA-induced tumors in these mice could result from impaired tumor antigen presentation via cross-dressing, cross-presentation or both processes. Addressing this question will require studying the growth of NLRC5-deficient and NLRC5-sufficient isogenic tumor cell lines in *Nlrc5^-/-^* and *Nlrc5^+/+^* hosts, or in mice lacking NLRC5 specifically in DCs.

Activated CD8^+^ T cells proliferate and differentiate into effector CTLs, enter circulation, traffic to the tumor, recognize tumor antigenic peptide:MHC-I complexes on cancer cells, and release cytolytic granules [62]. In addition to CD8^+^ T cells, natural killer (NK) cells also play a role in tumor immune surveillance, although they are not sufficient to control MCA-induced spontaneous tumors in *Rag1^-/-^* mice [70–72]. NK cells recognize cancer cells that lose the expression of classical MHC-Ia molecules, which engage NK cell inhibitory receptors, and cancer cells that express NK cell activating ligands, which include non-classical MHC-Ib molecules [73–77]. Tumors from *Nlrc5^-/-^*mice showed reduced expression of classical (H2-K1, H2-D1) and non-classical (H2-Q7, H-2-Q4) MHC-I molecules (Figure 7C). Earlier studies have shown that NLRC5 transactivated both MHC-Ia and MHC-Ib molecules [38,40,44,69], indicating a role for NLRC5 in activating both CTL and NK cell responses against tumors. A similar expression pattern of MHC-Ia and MHC-Ib molecules in *Nlrc5^-/-^* and *Rag^-/-^* tumors (Figure 7C) suggests their induction by ongoing immune responses, presumably via IFNγ and downstream induction of NLRC5. Granzymes are a key arsenal of cytolytic granules of CTLs and NK cells in killing cancer cells, of which GZMB is plays a major role [60,78,79]. Notably, GZMB expression that was markedly downregulated in *Rag^-/-^* tumors was only moderately reduced in *Nlrc5^-/-^* tumors (Figure 7A), suggesting its expression in other immune cells that may be activated in an NLRC5-independent fashion. Another notable difference related to CD8 receptor binding is the very low expression of FYN in *Nlrc5^-/-^* tumors. As FYN has been reported to attenuate T cell activation and commitment to effector cell differentiation [80,81], it raises the possibility that NLRC5 may have a role in alleviating this negative regulatory mechanism in CD8^+^ T cells. Besides CTLs and NK cells, γδ T cells may also contribute to elevated cytolytic responses in *Nlrc5^+/+^* tumors, as NLRC5 has been shown to promote killing by γδ T cells via induction of butyrophilin (BTN) family proteins BTN3A1-3 [82–84]. BTN proteins bound to altered self-proteins such as phospho-Ags in stressed and cancer cells are recognized by the γδ TCR in an MHC-independent manner [82].

Analysis of differentially expressed proteins in tumors from *Nlrc5^−/−^*, *Nlrc5^+/+^* and *Rag^-/-^* tumors revealed not only the enrichment of cytolytic immune cell responses in *Nlrc5^+/+^* tumors but also revealed suppression of humoral and innate immune responses, which are enriched in *Nlrc5^−/−^*tumors. This enrichment is particularly associated with IGHV peptides derived from antibodies, suggesting a skewed antigen reactivity of the tumor-associated B cell antibody repertoire. Tumor-associated B lymphocytes also actively participate in antitumor immune responses, however they are heterogeneous and may exert either anti-tumor or pro-tumor functions, and antibodies can cause aberrant autoimmune-like reactions in tumor microenvironment [85–87]. *Nlrc5^−/−^* tumors also displayed enrichment of GO terms related to proteolytic pathways, suggesting that loss of NLRC5-dependent classical MHC-I antigen processing pathway could trigger alternate antigen processing pathways [88]. The enrichment of distinct antibody responses and proteolytic pathways likely represent adaptations to compensate for the loss of NLRC5-dependent cytolytic immune effector cell functions. These adaptations may contribute to T cell infiltration and other immune effector cells giving rise to the necrotic areas in *Nlrc5^-/-^* tumors that were not present in *Rag1^−/−^* tumors. Evidently, these adaptations were insufficient to control tumor growth in *Nlrc5^-/-^* mice as their tumor incidence and survival were not significantly different from *Rag1^−/−^*mice.

Our findings show that NLRC5 is needed for robust immune surveillance against endogenously arising tumors. NLRC5 expression in tumors promote enrichment of proteins involved in activation of antigen processing and presentation, CD8 T cell receptor binding and granzyme-mediated cytolytic immune effector cell pathways. NLRC5 plays an indispensable role in tumor immunoediting. The loss of NLRC5-dependent adaptive tumor immune surveillance mechanisms promotes compensatory activation of innate and humoral immune response pathways, however, these pathways are not sufficient for cancer immune surveillance and cancer immunoediting.

## Acknowledgments

This research was funded by the Canadian Institutes of Health Research (CIHR) PJT-153174 to S.I. This research was enabled in part by support provided the team at the Centre de calcul scientifique de l’Université de Sherbrooke, Calcul Québec (calculquebec.ca) and the Digital Research Alliance of Canada (alliancecan.ca). CR-CHUS is an FRQS-funded research center.

## Figure Legends

**Supplementary Figure S1.**
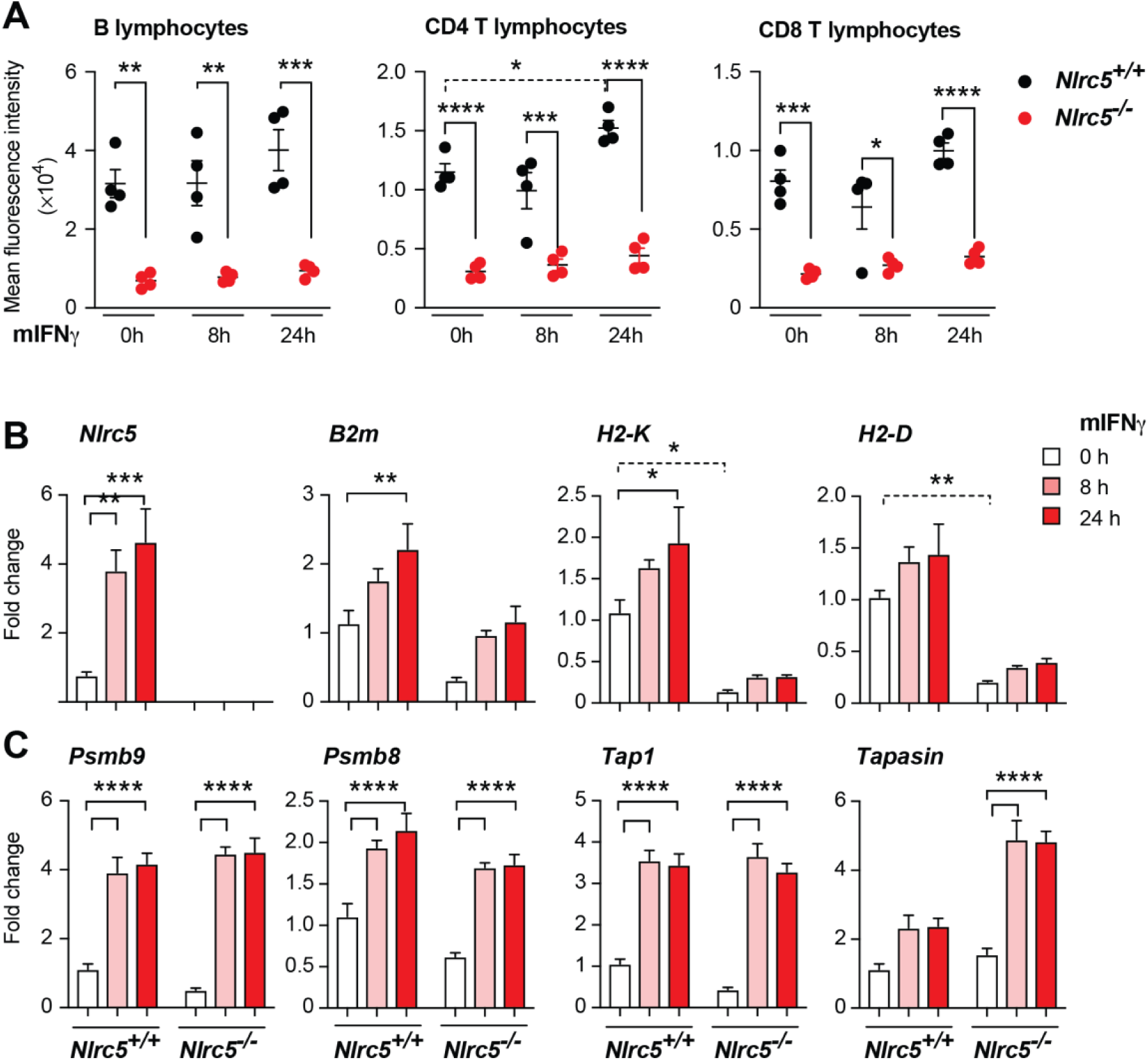
Validation of *Nlrc5^−/−^* mice. (A) Flow cytometry evaluation of MHC-I expression in CD45 gated B cells, CD4+ T cells and CD8+ T cells with and without and IFNγ stimulation (20 ng mL^−1^) for the indicated periods. (B, C) NLRC5 deficiency impairs the induction of MHC-I (B) and antigen processing machinery genes (C). Freshly isolated and IFNγ-stimulated (20ng/ml) splenocytes from *Nlrc5^-/-^* and *Nlrc5^+/+^* mice were evaluated for the expression of *Nlrc5, B2m*, *H2-K* and *H2-D* genes (B), and genes coding for LMP2 (*Psmb9*), LMP7 (*Psmb8*), TAP1 (*Tap1*) and Tapasin (*Tapbp*) (C) by RT-qPCR. The gene expression in each sample was normalized using the housekeeping gene β-actin to calculate the fold change following IFNγ stimulation. n=4 mice per group. Statistical significance using Turkey’s multiple comparison test. *p* values * ≤0.05, ** p≤0.01, *** p≤0.001, **** p≤0.0001.

## References

1. Hanahan, D.; Weinberg, R.A. Hallmarks of cancer: the next generation. Cell 2011, 144, 646–674; doi:10.1016/j.cell.2011.02.013.

2. Seliger, B.; Maeurer, M.J.; Ferrone, S. TAP off--tumors on. Immunol Today 1997, 18, 292–299; doi:10.1016/s0167-5699(97)01052-9.

3. Lampen, M.H.; van Hall, T. Strategies to counteract MHC-I defects in tumors. Curr Opin Immunol 2011, 23, 293–298; doi:10.1016/j.coi.2010.12.005.

4. Bukur, J.; Jasinski, S.; Seliger, B. The role of classical and non-classical HLA class I antigens in human tumors. Semin Cancer Biol 2012, 22, 350–358; doi:10.1016/j.semcancer.2012.03.003.

5. Dhatchinamoorthy, K.; Colbert, J.D.; Rock, K.L. Cancer Immune Evasion Through Loss of MHC Class I Antigen Presentation. Front Immunol 2021, 12, 636568; doi:10.3389/fimmu.2021.636568.

6. Campoli, M.; Chang, C.C.; Ferrone, S. HLA class I antigen loss, tumor immune escape and immune selection. Vaccine 2002, 20 <u>Suppl 4</u>, A40–45; doi:10.1016/s0264-410x(02)00386-9.

7. Cabrera, T.; Lara, E.; Romero, J.M.; Maleno, I.; Real, L.M.; Ruiz-Cabello, F.; Valero, P.; Camacho, F.M.; Garrido, F. HLA class I expression in metastatic melanoma correlates with tumor development during autologous vaccination. Cancer Immunol Immunother 2007, 56, 709–717; doi:10.1007/s00262-006-0226-7.

8. Carretero, R.; Romero, J.M.; Ruiz-Cabello, F.; Maleno, I.; Rodriguez, F.; Camacho, F.M.; Real, L.M.; Garrido, F.; Cabrera, T. Analysis of HLA class I expression in progressing and regressing metastatic melanoma lesions after immunotherapy. Immunogenetics 2008, 60, 439–447; doi:10.1007/s00251-008-0303-5.

9. Sokol, L.; Koelzer, V.H.; Rau, T.T.; Karamitopoulou, E.; Zlobec, I.; Lugli, A. Loss of tapasin correlates with diminished CD8(+) T-cell immunity and prognosis in colorectal cancer. J Transl Med 2015, 13, 279; doi:10.1186/s12967-015-0647-1.

10. Seliger, B. Molecular mechanisms of MHC class I abnormalities and APM components in human tumors. Cancer Immunol Immunother 2008, 57, 1719–1726; doi:10.1007/s00262-008-0515-4.

11. Garrido, F.; Cabrera, T.; Aptsiauri, N. “Hard” and “soft” lesions underlying the HLA class I alterations in cancer cells: implications for immunotherapy. Int J Cancer 2010, 127, 249–256; doi:10.1002/ijc.25270.

12. Garrido, F.; Algarra, I.; Garcia-Lora, A.M. The escape of cancer from T lymphocytes: immunoselection of MHC class I loss variants harboring structural-irreversible “hard” lesions. Cancer Immunol Immunother 2010, 59, 1601–1606; doi:10.1007/s00262-010-0893-2.

13. Kloetzel, P.M. Antigen processing by the proteasome. Nat Rev Mol Cell Biol 2001, 2, 179–187; doi:10.1038/35056572.

14. Tanaka, K. The proteasome: overview of structure and functions. Proc Jpn Acad Ser B Phys Biol Sci 2009, 85, 12–36; doi:10.2183/pjab.85.12.

15. Amm, I.; Sommer, T.; Wolf, D.H. Protein quality control and elimination of protein waste: the role of the ubiquitin-proteasome system. Biochim Biophys Acta 2014, 1843, 182–196; doi:10.1016/j.bbamcr.2013.06.031.

16. Yewdell, J.W.; Reits, E.; Neefjes, J. Making sense of mass destruction: quantitating MHC class I antigen presentation. Nat Rev Immunol 2003, 3, 952–961; doi:10.1038/nri1250.

17. Eggensperger, S.; Tampe, R. The transporter associated with antigen processing: a key player in adaptive immunity. Biol Chem 2015, 396, 1059–1072; doi:10.1515/hsz-2014-0320.

18. Hulpke, S.; Tampe, R. The MHC I loading complex: a multitasking machinery in adaptive immunity. Trends Biochem Sci 2013, 38, 412–420; doi:10.1016/j.tibs.2013.06.003.

19. Benitez, R.; Godelaine, D.; Lopez-Nevot, M.A.; Brasseur, F.; Jimenez, P.; Marchand, M.; Oliva, M.R.; van Baren, N.; Cabrera, T.; Andry, G.;, et al. Mutations of the beta2-microglobulin gene result in a lack of HLA class I molecules on melanoma cells of two patients immunized with MAGE peptides. Tissue Antigens 1998, 52, 520–529; doi:10.1111/j.1399-0039.1998.tb03082.x.

20. Hicklin, D.J.; Marincola, F.M.; Ferrone, S. HLA class I antigen downregulation in human cancers: T-cell immunotherapy revives an old story. Mol Med Today 1999, 5, 178–186; doi:10.1016/s1357-4310(99)01451-3.

21. Leone, P.; Shin, E.C.; Perosa, F.; Vacca, A.; Dammacco, F.; Racanelli, V. MHC class I antigen processing and presenting machinery: organization, function, and defects in tumor cells. J Natl Cancer Inst 2013, 105, 1172–1187; doi:10.1093/jnci/djt184.

22. Chen, H.L.; Gabrilovich, D.; Tampe, R.; Girgis, K.R.; Nadaf, S.; Carbone, D.P. A functionally defective allele of TAP1 results in loss of MHC class I antigen presentation in a human lung cancer. Nat Genet 1996, 13, 210–213; doi:10.1038/ng0696-210.

23. Seliger, B.; Ritz, U.; Abele, R.; Bock, M.; Tampe, R.; Sutter, G.; Drexler, I.; Huber, C.; Ferrone, S. Immune escape of melanoma: first evidence of structural alterations in two distinct components of the MHC class I antigen processing pathway. Cancer Res 2001, 61, 8647–8650.

24. Shankaran, V.; Ikeda, H.; Bruce, A.T.; White, J.M.; Swanson, P.E.; Old, L.J.; Schreiber, R.D. IFNgamma and lymphocytes prevent primary tumour development and shape tumour immunogenicity. Nature 2001, 410, 1107–1111; doi:10.1038/35074122.

25. Dunn, G.P.; Old, L.J.; Schreiber, R.D. The immunobiology of cancer immunosurveillance and immunoediting. Immunity 2004, 21, 137–148.

26. Magalhaes, J.G.; Sorbara, M.T.; Girardin, S.E.; Philpott, D.J. What is new with Nods? Curr Opin Immunol 2011, 23, 29–34; doi:10.1016/j.coi.2010.12.003.

27. Motta, V.; Soares, F.; Sun, T.; Philpott, D.J. NOD-like receptors: versatile cytosolic sentinels. Physiol Rev 2015, 95, 149–178; doi:10.1152/physrev.00009.2014.

28. Cui, J.; Zhu, L.; Xia, X.; Wang, H.Y.; Legras, X.; Hong, J.; Ji, J.; Shen, P.; Zheng, S.; Chen, Z.J.;, et al. NLRC5 negatively regulates the NF-kappaB and type I interferon signaling pathways. Cell 2010, 141, 483–496; doi:10.1016/j.cell.2010.03.040.

29. Benko, S.; Magalhaes, J.G.; Philpott, D.J.; Girardin, S.E. NLRC5 limits the activation of inflammatory pathways. J Immunol 2010, 185, 1681–1691; doi:10.4049/jimmunol.0903900.

30. Neerincx, A.; Lautz, K.; Menning, M.; Kremmer, E.; Zigrino, P.; Hosel, M.; Buning, H.; Schwarzenbacher, R.; Kufer, T.A. A role for the human nucleotide-binding domain, leucine-rich repeat-containing family member NLRC5 in antiviral responses. J Biol Chem 2010, 285, 26223–26232; doi:10.1074/jbc.M110.109736.

31. Meng, Q.; Cai, C.; Sun, T.; Wang, Q.; Xie, W.; Wang, R.; Cui, J. Reversible ubiquitination shapes NLRC5 function and modulates NF-kappaB activation switch. J Cell Biol 2015, 211, 1025–1040; doi:10.1083/jcb.201505091.

32. Hu, Y. A feedforward loop of NLRC5 (de)ubiquitination keeps IKK-NF-kappaB in check. J Cell Biol 2015, 211, 941–943; doi:10.1083/jcb.201511039.

33. Meissner, T.B.; Li, A.; Biswas, A.; Lee, K.H.; Liu, Y.J.; Bayir, E.; Iliopoulos, D.; van den Elsen, P.J.; Kobayashi, K.S. NLR family member NLRC5 is a transcriptional regulator of MHC class I genes. Proc Natl Acad Sci U S A 2010, 107, 13794–13799; doi:10.1073/pnas.1008684107.

34. Kobayashi, K.S.; van den Elsen, P.J. NLRC5: a key regulator of MHC class I-dependent immune responses. Nat Rev Immunol 2012, 12, 813–820; doi:10.1038/nri3339.

35. Meissner, T.B.; Liu, Y.J.; Lee, K.H.; Li, A.; Biswas, A.; van Eggermond, M.C.; van den Elsen, P.J.; Kobayashi, K.S. NLRC5 cooperates with the RFX transcription factor complex to induce MHC class I gene expression. J Immunol 2012, 188, 4951–4958; doi:10.4049/jimmunol.1103160.

36. Neerincx, A.; Rodriguez, G.M.; Steimle, V.; Kufer, T.A. NLRC5 controls basal MHC class I gene expression in an MHC enhanceosome-dependent manner. J Immunol 2012, 188, 4940–4950; doi:10.4049/jimmunol.1103136.

37. Meissner, T.B.; Li, A.; Liu, Y.J.; Gagnon, E.; Kobayashi, K.S. The nucleotide-binding domain of NLRC5 is critical for nuclear import and transactivation activity. Biochem Biophys Res Commun 2012, 418, 786–791; doi:10.1016/j.bbrc.2012.01.104.

38. Robbins, G.R.; Truax, A.D.; Davis, B.K.; Zhang, L.; Brickey, W.J.; Ting, J.P. Regulation of class I major histocompatibility complex (MHC) by nucleotide-binding domain, leucine-rich repeat-containing (NLR) proteins. J Biol Chem 2012, 287, 24294–24303; doi:10.1074/jbc.M112.364604.

39. Staehli, F.; Ludigs, K.; Heinz, L.X.; Seguin-Estevez, Q.; Ferrero, I.; Braun, M.; Schroder, K.; Rebsamen, M.; Tardivel, A.; Mattmann, C.;, et al. NLRC5 deficiency selectively impairs MHC class I-dependent lymphocyte killing by cytotoxic T cells. J Immunol 2012, 188, 3820–3828; doi:10.4049/jimmunol.1102671.

40. Ludigs, K.; Jandus, C.; Utzschneider, D.T.; Staehli, F.; Bessoles, S.; Dang, A.T.; Rota, G.; Castro, W.; Zehn, D.; Vivier, E.;, et al. NLRC5 shields T lymphocytes from NK-cell-mediated elimination under inflammatory conditions. Nat Commun 2016, 7, 10554; doi:10.1038/ncomms10554.

41. Sun, T.; Ferrero, R.L.; Girardin, S.E.; Gommerman, J.L.; Philpott, D.J. NLRC5 deficiency has a moderate impact on immunodominant CD8(+) T cell responses during rotavirus infection of adult mice. Immunol Cell Biol 2019; doi:10.1111/imcb.12244.

42. Vugmeyster, Y.; Glas, R.; Perarnau, B.; Lemonnier, F.A.; Eisen, H.; Ploegh, H. Major histocompatibility complex (MHC) class I KbDb -/- deficient mice possess functional CD8+ T cells and natural killer cells. Proc Natl Acad Sci U S A 1998, 95, 12492–12497; doi:10.1073/pnas.95.21.12492.

43. Grusby, M.J.; Auchincloss, H., Jr.; Lee, R.; Johnson, R.S.; Spencer, J.P.; Zijlstra, M.; Jaenisch, R.; Papaioannou, V.E.; Glimcher, L.H. Mice lacking major histocompatibility complex class I and class II molecules. Proc Natl Acad Sci U S A 1993, 90, 3913–3917; doi:10.1073/pnas.90.9.3913.

44. Yao, Y.; Wang, Y.; Chen, F.; Huang, Y.; Zhu, S.; Leng, Q.; Wang, H.; Shi, Y.; Qian, Y. NLRC5 regulates MHC class I antigen presentation in host defense against intracellular pathogens. Cell Res 2012, 22, 836–847; doi:10.1038/cr.2012.56.

45. Benhammadi, M.; Mathe, J.; Dumont-Lagace, M.; Kobayashi, K.S.; Gaboury, L.; Brochu, S.; Perreault, C. IFN-lambda Enhances Constitutive Expression of MHC Class I Molecules on Thymic Epithelial Cells. J Immunol 2020, 205, 1268–1280; doi:10.4049/jimmunol.2000225.

46. Yoshihama, S.; Roszik, J.; Downs, I.; Meissner, T.B.; Vijayan, S.; Chapuy, B.; Sidiq, T.; Shipp, M.A.; Lizee, G.A.; Kobayashi, K.S. NLRC5/MHC class I transactivator is a target for immune evasion in cancer. Proc Natl Acad Sci U S A 2016, 113, 5999–6004; doi:10.1073/pnas.1602069113.

47. Yoshihama, S.; Cho, S.X.; Yeung, J.; Pan, X.; Lizee, G.; Konganti, K.; Johnson, V.E.; Kobayashi, K.S. NLRC5/CITA expression correlates with efficient response to checkpoint blockade immunotherapy. Sci Rep 2021, 11, 3258; doi:10.1038/s41598-021-82729-9.

48. Rodriguez, G.M.; Bobbala, D.; Serrano, D.; Mayhue, M.; Champagne, A.; Saucier, C.; Steimle, V.; Kufer, T.A.; Menendez, A.; Ramanathan, S.;, et al. NLRC5 elicits antitumor immunity by enhancing processing and presentation of tumor antigens to CD8(+) T lymphocytes. Oncoimmunology 2016, 5, e1151593; doi:10.1080/2162402X.2016.1151593.

49. Rodriguez, G.M.; Yakubovich, E.; Murshed, H.; Maranda, V.; Galpin, K.J.C.; Cudmore, A.; Hanna, A.M.R.; Macdonald, E.; Ramesh, S.; Garson, K.;, et al. NLRC5 overexpression in ovarian tumors remodels the tumor microenvironment and increases T-cell reactivity toward autologous tumor-associated antigens. Front Immunol 2023, 14, 1295208; doi:10.3389/fimmu.2023.1295208.

50. Yoshihama, S.; Vijayan, S.; Sidiq, T.; Kobayashi, K.S. NLRC5/CITA: A Key Player in Cancer Immune Surveillance. Trends Cancer 2017, 3, 28–38; doi:10.1016/j.trecan.2016.12.003.

51. Quenum, A.J.I.; Shukla, A.; Rexhepi, F.; Cloutier, M.; Ghosh, A.; Kufer, T.A.; Ramanathan, S.; Ilangumaran, S. NLRC5 Deficiency Deregulates Hepatic Inflammatory Response but Does Not Aggravate Carbon Tetrachloride-Induced Liver Fibrosis. Front Immunol 2021, 12, 749646; doi:10.3389/fimmu.2021.749646.

52. Schreiber, T.H.; Podack, E.R. A critical analysis of the tumour immunosurveillance controversy for 3-MCA-induced sarcomas. Br J Cancer 2009, 101, 381–386; doi:10.1038/sj.bjc.6605198.

53. Stagg, J.; Beavis, P.A.; Divisekera, U.; Liu, M.C.; Moller, A.; Darcy, P.K.; Smyth, M.J. CD73-deficient mice are resistant to carcinogenesis. Cancer Res 2012, 72, 2190–2196; doi:10.1158/0008-5472.CAN-12-0420.

54. Demichev, V.; Messner, C.B.; Vernardis, S.I.; Lilley, K.S.; Ralser, M. DIA-NN: neural networks and interference correction enable deep proteome coverage in high throughput. Nat Methods 2020, 17, 41–44; doi:10.1038/s41592-019-0638-x.

55. https://hub.docker.com/layers/biocontainers/diann/v1.8.1_cv1/images/sha256-c37bb6b4baa8bcc1552b8ddc103fe5e43731832feb2ed739e4b6c2cb9cf78471.

56. Hsiao, Y.; Zhang, H.; Li, G.X.; Deng, Y.; Yu, F.; Valipour Kahrood, H.; Steele, J.R.; Schittenhelm, R.B.; Nesvizhskii, A.I. Analysis and Visualization of Quantitative Proteomics Data Using FragPipe-Analyst. J Proteome Res 2024, 23, 4303–4315; doi:10.1021/acs.jproteome.4c00294.

57. Bardou, P.; Mariette, J.; Escudie, F.; Djemiel, C.; Klopp, C. jvenn: an interactive Venn diagram viewer. BMC Bioinformatics 2014, 15, 293; doi:10.1186/1471-2105-15-293.

58. Wang, X.; Terfve, C.; Rose, J.C.; Markowetz, F. HTSanalyzeR: an R/Bioconductor package for integrated network analysis of high-throughput screens. Bioinformatics 2011, 27, 879–880; doi:10.1093/bioinformatics/btr028.

59. Shukla, A.; Cloutier, M.; Appiya Santharam, M.; Ramanathan, S.; Ilangumaran, S. The MHC Class-I Transactivator NLRC5: Implications to Cancer Immunology and Potential Applications to Cancer Immunotherapy. Int J Mol Sci 2021, 22; doi:10.3390/ijms22041964.

60. Trapani, J.A. Granzymes: a family of lymphocyte granule serine proteases. Genome Biol 2001, 2, REVIEWS3014; doi:10.1186/gb-2001-2-12-reviews3014.

61. Santharam, M.A.; Shukla, A.; Levesque, D.; Kufer, T.A.; Boisvert, F.M.; Ramanathan, S.; Ilangumaran, S. NLRC5-CIITA Fusion Protein as an Effective Inducer of MHC-I Expression and Antitumor Immunity. Int J Mol Sci 2023, 24; doi:10.3390/ijms24087206.

62. Chen, D.S.; Mellman, I. Oncology meets immunology: the cancer-immunity cycle. Immunity 2013, 39, 1–10; doi:10.1016/j.immuni.2013.07.012.

63. Jhunjhunwala, S.; Hammer, C.; Delamarre, L. Antigen presentation in cancer: insights into tumour immunogenicity and immune evasion. Nat Rev Cancer 2021, 21, 298–312; doi:10.1038/s41568-021-00339-z.

64. Hoffmann, T.K.; Meidenbauer, N.; Dworacki, G.; Kanaya, H.; Whiteside, T.L. Generation of tumor-specific T-lymphocytes by cross-priming with human dendritic cells ingesting apoptotic tumor cells. Cancer Res 2000, 60, 3542–3549.

65. Sanchez-Paulete, A.R.; Teijeira, A.; Cueto, F.J.; Garasa, S.; Perez-Gracia, J.L.; Sanchez-Arraez, A.; Sancho, D.; Melero, I. Antigen cross-presentation and T-cell cross-priming in cancer immunology and immunotherapy. Ann Oncol 2017, 28, xii74; doi:10.1093/annonc/mdx727.

66. Nakayama, M. Antigen Presentation by MHC-Dressed Cells. Front Immunol 2014, 5, 672; doi:10.3389/fimmu.2014.00672.

67. Das Mohapatra, A.; Tirrell, I.; Benechet, A.P.; Pattnayak, S.; Khanna, K.M.; Srivastava, P.K. Cross-dressing of CD8alpha(+) Dendritic Cells with Antigens from Live Mouse Tumor Cells Is a Major Mechanism of Cross-priming. Cancer Immunol Res 2020, 8, 1287–1299; doi:10.1158/2326-6066.CIR-20-0248.

68. MacNabb, B.W.; Tumuluru, S.; Chen, X.; Godfrey, J.; Kasal, D.N.; Yu, J.; Jongsma, M.L.M.; Spaapen, R.M.; Kline, D.E.; Kline, J. Dendritic cells can prime anti-tumor CD8(+) T cell responses through major histocompatibility complex cross-dressing. Immunity 2022, 55, 982–997 e988; doi:10.1016/j.immuni.2022.04.016.

69. Biswas, A.; Meissner, T.B.; Kawai, T.; Kobayashi, K.S. Cutting edge: impaired MHC class I expression in mice deficient for Nlrc5/class I transactivator. J Immunol 2012, 189, 516–520; doi:10.4049/jimmunol.1200064.

70. Morvan, M.G.; Lanier, L.L. NK cells and cancer: you can teach innate cells new tricks. Nat Rev Cancer 2016, 16, 7–19; doi:10.1038/nrc.2015.5.

71. Meza Guzman, L.G.; Keating, N.; Nicholson, S.E. Natural Killer Cells: Tumor Surveillance and Signaling. Cancers (Basel) 2020, 12; doi:10.3390/cancers12040952.

72. Smyth, M.J.; Swann, J.; Cretney, E.; Zerafa, N.; Yokoyama, W.M.; Hayakawa, Y. NKG2D function protects the host from tumor initiation. J Exp Med 2005, 202, 583–588; doi:10.1084/jem.20050994.

73. Vivier, E.; Tomasello, E.; Baratin, M.; Walzer, T.; Ugolini, S. Functions of natural killer cells. Nat Immunol 2008, 9, 503–510; doi:10.1038/ni1582.

74. Long, E.O.; Kim, H.S.; Liu, D.; Peterson, M.E.; Rajagopalan, S. Controlling natural killer cell responses: integration of signals for activation and inhibition. Annu Rev Immunol 2013, 31, 227–258; doi:10.1146/annurev-immunol-020711-075005.

75. Kirkham, C.L.; Carlyle, J.R. Complexity and Diversity of the NKR-P1:Clr (Klrb1:Clec2) Recognition Systems. Front Immunol 2014, 5, 214; doi:10.3389/fimmu.2014.00214.

76. Paul, S.; Lal, G. The Molecular Mechanism of Natural Killer Cells Function and Its Importance in Cancer Immunotherapy. Front Immunol 2017, 8, 1124; doi:10.3389/fimmu.2017.01124.

77. Wolf, N.K.; Kissiov, D.U.; Raulet, D.H. Roles of natural killer cells in immunity to cancer, and applications to immunotherapy. Nat Rev Immunol 2023, 23, 90–105; doi:10.1038/s41577-022-00732-1.

78. Revell, P.A.; Grossman, W.J.; Thomas, D.A.; Cao, X.; Behl, R.; Ratner, J.A.; Lu, Z.H.; Ley, T.J. Granzyme B and the downstream granzymes C and/or F are important for cytotoxic lymphocyte functions. J Immunol 2005, 174, 2124–2131; doi:10.4049/jimmunol.174.4.2124.

79 Cullen, S.P.; Brunet, M.; Martin, S.J. Granzymes in cancer and immunity. Cell Death Differ 2010, 17, 616–623; doi:10.1038/cdd.2009.206.

80. Filby, A.; Seddon, B.; Kleczkowska, J.; Salmond, R.; Tomlinson, P.; Smida, M.; Lindquist, J.A.; Schraven, B.; Zamoyska, R. Fyn regulates the duration of TCR engagement needed for commitment to effector function. J Immunol 2007, 179, 4635–4644; doi:10.4049/jimmunol.179.7.4635.

81. Borger, J.G.; Filby, A.; Zamoyska, R. Differential polarization of C-terminal Src kinase between naive and antigen-experienced CD8+ T cells. J Immunol 2013, 190, 3089–3099; doi:10.4049/jimmunol.1202408.

82. Silva-Santos, B.; Mensurado, S.; Coffelt, S.B. gammadelta T cells: pleiotropic immune effectors with therapeutic potential in cancer. Nat Rev Cancer 2019, 19, 392–404; doi:10.1038/s41568-019-0153-5.

83. de Vries, N.L.; van de Haar, J.; Veninga, V.; Chalabi, M.; Ijsselsteijn, M.E.; van der Ploeg, M.; van den Bulk, J.; Ruano, D.; van den Berg, J.G.; Haanen, J.B.;, et al. gammadelta T cells are effectors of immunotherapy in cancers with HLA class I defects. Nature 2023, 613, 743–750; doi:10.1038/s41586-022-05593-1.

84. Dang, A.T.; Strietz, J.; Zenobi, A.; Khameneh, H.J.; Brandl, S.M.; Lozza, L.; Conradt, G.; Kaufmann, S.H.E.; Reith, W.; Kwee, I.;, et al. NLRC5 promotes transcription of BTN3A1-3 genes and Vgamma9Vdelta2 T cell-mediated killing. iScience 2021, 24, 101900; doi:10.1016/j.isci.2020.101900.

85. Tsou, P.; Katayama, H.; Ostrin, E.J.; Hanash, S.M. The Emerging Role of B Cells in Tumor Immunity. Cancer Res 2016, 76, 5597–5601; doi:10.1158/0008-5472.CAN-16-0431.

86. Qin, Y.; Lu, F.; Lyu, K.; Chang, A.E.; Li, Q. Emerging concepts regarding pro- and anti tumor properties of B cells in tumor immunity. Front Immunol 2022, 13, 881427; doi:10.3389/fimmu.2022.881427.

86. Crescioli, S.; Correa, I.; Ng, J.; Willsmore, Z.N.; Laddach, R.; Chenoweth, A.; Chauhan, J.; Di Meo, A.; Stewart, A.; Kalliolia, E.;, et al. B cell profiles, antibody repertoire and reactivity reveal dysregulated responses with autoimmune features in melanoma. Nat Commun 2023, 14, 3378; doi:10.1038/s41467-023-39042-y.

87. Cruz, F.M.; Orellano, L.A.A.; Chan, A.; Rock, K.L. Alternate MHC I Antigen Presentation Pathways Allow CD8+ T-cell Recognition and Killing of Cancer Cells in the Absence of beta2M or TAP. Cancer Immunol Res 2025, 13, 98–108; doi:10.1158/2326-6066.CIR-24-0320.

